# Molecular mechanisms of inorganic-phosphate release from the core and barbed end of actin filaments

**DOI:** 10.1101/2023.03.25.534205

**Authors:** Wout Oosterheert, Florian E.C. Blanc, Ankit Roy, Alexander Belyy, Oliver Hofnagel, Gerhard Hummer, Peter Bieling, Stefan Raunser

## Abstract

The release of inorganic phosphate (P_i_) from actin filaments constitutes a key step in their regulated turnover, which is fundamental to many cellular functions. However, the molecular mechanisms underlying P_i_ release from both the core and barbed end of actin filaments remain unclear. Here, we combine cryo-EM with molecular dynamics simulations and *in vitro* reconstitution to demonstrate how actin releases P_i_ through a ‘molecular backdoor’. While constantly open at the barbed end, the backdoor is predominantly closed in filament-core subunits and only opens transiently through concerted backbone movements and rotameric rearrangements of residues close to the nucleotide binding pocket. This mechanism explains why P_i_ escapes rapidly from the filament end and yet slowly from internal actin subunits. In an actin variant associated with nemaline myopathy, the backdoor is predominantly open in filament-core subunits, resulting in greatly accelerated P_i_ release after polymerization and filaments with drastically shortened ADP-P_i_ caps. This demonstrates that the P_i_ release rate from F-actin is controlled by steric hindrance through the backdoor rather than by the disruption of the ionic bond between P_i_ and Mg^2+^ at the nucleotide-binding site. Our results provide the molecular basis for P_i_ release from actin and exemplify how a single, disease-linked point mutation distorts the nucleotide state distribution and atomic structure of the actin filament.

## Introduction

The dynamic turnover of actin filaments (F-actin) controls the shape and movement of eukaryotic cells and is driven by changes in the molecular identity of the adenine-nucleotide bound to actin^1–3^. While monomeric actin (G-actin) displays only very weak hydrolysis activity towards ATP^4^, actin polymerization results in the flattening of the protein and a rearrangement of amino acids and water molecules near the nucleotide-binding site^5–8^. Accordingly, the ATPase activity of F-actin increases about 42,000 fold and hydrolysis takes place within seconds of filament formation (rate 0.3 s^-1^) (ref.^9^). Because the release of the cleaved inorganic phosphate (P_i_) occurs at a much slower rate than ATP hydrolysis^10^, the cap of a growing filament is generally rich in ADP-P_i_-bound subunits^11^, whereas ‘aged’ F-actin primarily adopts the ADP-bound state. *In vivo*, these changes in the nucleotide state are sensed by a variety of actin-binding proteins (ABPs)^12–14^. For instance, ADF/cofilin family proteins efficiently bind and sever the ADP-bound state of the filament, but bind only weakly to ADP-P_i_-F-actin^15–17^. Thus, the release of P_i_ from the F-actin interior represents a crucial step in the control of filament turnover.

For a long time, it was debated whether or not the P_i_ release rate of a given F-actin subunit depends on the nucleotide state of its neighboring subunits^10, 18, 19^. Finally, single-filament experiments provided experimental evidence that P_i_ release from the F-actin core is a stochastic process; each ADP-P_i_-subunit in the filament releases its bound phosphate molecule with equal probability at a rate of 0.002 to 0.007 s^-1^, which corresponds to a half-time τ of several minutes^10, 20–24^. Interestingly, under depolymerization conditions, actin subunits at the barbed end release P_i_ more than 300-fold faster (∼2 s^-1^) than those that reside in the filament core, even though the equilibrium constants of P_i_ binding to barbed end and filament core are essentially the same^20, 25^. These observations therefore imply that P_i_ binding is also much faster at the barbed end of F-actin^25^.

At the atomic level, P_i_ release from actin has been investigated by pioneering molecular dynamics (MD) simulation studies in the late 1990s (refs.^26, 27^), which suggested that the disruption of the ionic bond between P_i_ and the nucleotide-associated divalent cation (Mg^2+^ or Ca^2+^) could represent the rate limiting step for P_i_ release, because actin rearrangements were not required for P_i_ to exit. These studies furthermore predicted that P_i_ escapes through an open ‘backdoor’ in the actin molecule, with residues arginine-177 (R177) and methylated histidine-73 (H73) as potential mediators of P_i_ release. However, a central limitation of the simulations was that they were performed on a G-actin structure^28^, because no high-resolution F-actin structures were available at the time. In 2015, the first sub-4 Å cryo-EM structures of F-actin revealed that within the flattened conformation of the filament, P_i_ cannot freely diffuse out of the F-actin interior^29^; R177 participates in a hydrogen bonding network with the side chain of N111 and the backbones of H73 and G74, defining a closed backdoor conformation in F-actin. Accordingly, recently published cryo-EM structures of F-actin bound to ADP-P_i_ and ADP at resolutions beyond 2.5 Å showed a closed backdoor in both the pre- and post-release states^8, 30^. These data therefore suggest that P_i_ release from the filament interior occurs through a transient, high-energy state of F-actin that requires substantial rearrangements, but is difficult to capture with static imaging techniques such as cryo-EM. In a more recently proposed model for P_i_-release, a rotameric switch of residue S14 from a hydrogen-bond interaction with the backbone amide of G74 to the one of G158 would enable P_i_ to approach the R177-N111 backdoor and egress^31^. However, in the absence of experimental validation, the molecular principles that underlie P_i_ release from F-actin remain elusive, and further evidence is required to determine whether the phosphate molecule exits the filament interior through the postulated backdoor or through potential other egress routes. Additionally, the structural basis for the orders-of-magnitude faster P_i_ release from actin subunits at the barbed end compared to those that reside in the filament core is still unknown.

Here, we uncover that P_i_ release from subunits at the filament core and at the barbed end occurs through a common backdoor pathway. For internal subunits, the backdoor is predominantly closed and its transient opening is kinetically limited, whereas at the barbed end, the backdoor is open and P_i_ is capable of escaping from the ultimate subunit without large protein rearrangements. Strikingly, we also characterize an actin disease variant (N111S) that adopts an open backdoor arrangement in internal F-actin subunits and, hence, releases P_i_ without considerable delay. Our results provide a detailed molecular description of P_i_ release from F-actin and highlight a general approach of studying the mechanism of disease-linked actin variants.

## Results

### Structure of the F-actin barbed end reveals an open backdoor in the ultimate subunit

To elucidate how the conformation of the F-actin barbed end allows for much faster P_i_ release kinetics during filament depolymerization, we first aimed to obtain structural information on the barbed end by single-particle cryo-EM. Through a novel experimental approach, in which we polymerized free G-actin (native bovine β/γ-actin) in the presence of G-actin:DNase I complex, the FH2 domain of formin mDia1 (mDia1_FH2_) and the filament-stabilizing toxin phalloidin (see Methods for details), we reproducibly formed short (∼50 – 150 nm length) filaments and obtained a cryo-EM structure of the barbed-end at 3.6-Å resolution (Fig. 1a, b Supplementary Fig. 1, Supplementary Table 1, Supplementary Video 1). Importantly, we did not find any evidence for an ABP remaining bound to the barbed end, defining our structure as the undecorated barbed end of F-actin.

**Fig. 1.**
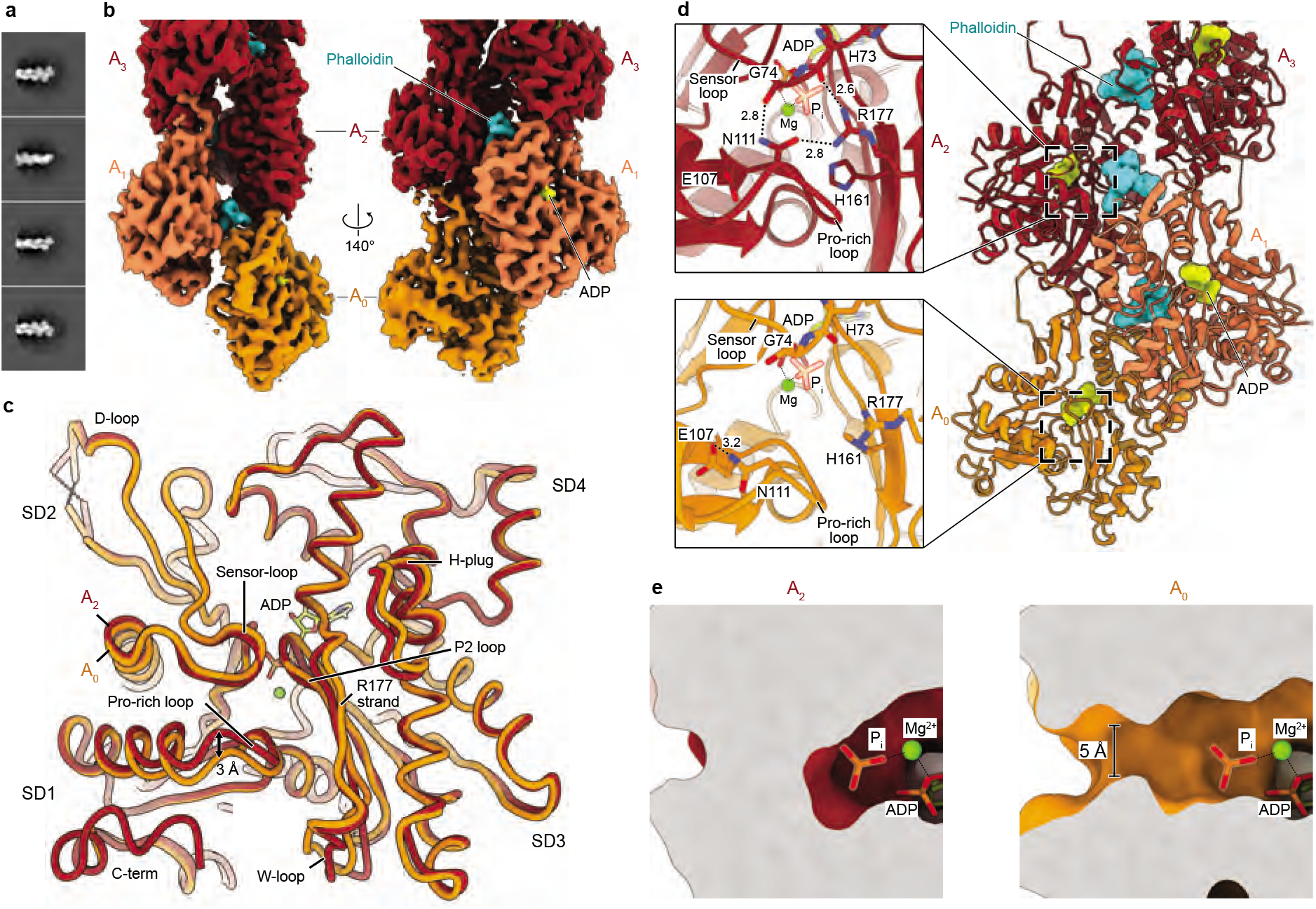
Cryo-EM structure of the phalloidin-bound barbed end of F-actin. **a** Exemplary 2D class averages of the barbed-end particles. The particle boxes are 465 × 465 Å^2^. Additional 2D class averages are depicted in Supplementary Fig. 1b. **b** Cryo-EM density map of the barbed end at 3.6-Å resolution, shown in two orientations. Actin subunits with complete inter-subunit contacts (A_2_ and above) are colored dark-red, the penultimate subunit (A_1_) is colored orange-red and the ultimate subunit (A_0_) is depicted in orange. Phalloidin (cyan) and ADP (yellow) are annotated. **c** Superimposition between the A_2_ and A_0_ subunits at the barbed end. Actin subdomains (SD1-4) and the regions that are different between the two subunits are annotated. P2 loop: residues 154-161. R177-strand: residues 176-178. **d** Right: Molecular model of the F-actin barbed end. The images on the left depicts a zoom-in on the R177-N111 backdoor as seen from the F-actin exterior. Amino acids that form the backdoor are annotated. Although the subunits adopt the ADP state, P_i_ from pdb 8a2s (F-actin in the Mg^2+^-ADP-P_i_ state) is fitted and shown semi-transparently to emphasize the P_i_-binding site. **e** Slice through the structures of the A_2_ and A_0_ subunits near the nucleotide-binding site, shown in surface representation. Similar to panel (**d**), P_i_ from pdb 8a2s is shown to emphasize the P_i_-binding site.

The structure reveals the hallmark actin filament architecture of a double-stranded helix with a right-handed twist (Fig. 1b, d). All actin subunits in our structure adopt the aged ADP- nucleotide state (Supplementary Fig. 2a, b). The arrangements of F-actin subunits that make all available inter-subunit contacts within the filament (A_2_ and above), as well as that of the penultimate subunit A_1_, are essentially the same as in previously solved F-actin structures (Supplementary Fig. 3, Cα rmsd <0.5 Å with pdb 8a2t)^8^. Accordingly, in subunits A_1_ and above, the predicted R177-N111 backdoor is closed and the P_i_-binding site is shielded from the filament exterior (Fig. 1d, Supplementary Fig. 3e). Although the overall arrangement of the ultimate (A_0_) subunit is also similar (Cα rmsd of 0.9 Å between subunits A_0_ and A_2_) and it remains in a flattened conformation (Supplementary Fig. 3c), we observed several differences when compared to the internal subunit A_2_, including small rearrangements of the W-loop (residues 165 – 172), the hydrophobic plug region (residues 264 – 273) and a disordered F-actin C-terminus (residues 363 – 375) (Fig. 1c). However, the most striking rearrangement is the downward displacement of the Pro-rich loop (residues 107 – 112) by ∼3 Å (Fig. 1c), which – unlike in internal actin subunits – it is not stabilized by subdomain 4 (SD4) of an adjacent subunit (Supplementary Fig. 2d). Hence, N111 loses its hydrogen bonds with G74 of the sensor loop (residues 70 – 77) and SD3-residue R177, and instead interacts within its own local loop with E107 (Fig. 1d, Supplementary Fig. 3f). Interestingly, this E107-N111 interaction is commonly observed in G-actin structures (Supplementary Fig. 3f). Strikingly, residue H161, which flips rotameric position during the G- to F-actin transition and is important for ATP hydrolysis in F-actin^7, 8^, also adopts its G-actin-like rotameric position in the A_0_ subunit and points towards R177. As a result, R177 can no longer interact with the sensor loop (Fig. 1d, Supplementary Fig. 3f). When taken together, the hydrogen-bonding network formed by R177, N111, H73 and G74 is fully abolished, which opens a ∼5 Å diameter hole in the structure that connects the internal nucleotide-binding site to the filament exterior (Fig. 1e, Supplementary Video 1). This defines the predicted^27^ P_i_ release backdoor as open. Thus, our structure provides evidence that, under depolymerization conditions, P_i_ can dissociate from the nucleotide-binding site at the barbed end without large protein rearrangements, thereby revealing the structural basis for the orders-of-magnitude faster P_i_ release rates from the ultimate barbed end subunit compared to those within the filament core.

### Two plausible P_i_ egress routes from the F-actin core

The barbed end structure revealed a P_i_ release pathway similar to the path predicted in the MD simulation study by Wriggers and Schulten from 1999 (ref.^27^). Accordingly, when we performed MD simulations on P_i_ release from barbed end subunit A_0_, we observed that virtually all P_i_ escape events occurred through the open R177-N111 backdoor (Supplementary Fig. 4a, b). However, we reasoned that for actin subunits in the filament core, which make all available inter-subunit contacts and release P_i_ at much slower rates, other P_i_ exit routes might exist. We therefore set out to develop an MD protocol to investigate the P_i_ release mechanism from the F-actin filament core, using our recently reported ∼2.2-Å structure of F-actin in the Mg^2+^-ADP- P_i_ state^8^ as high quality starting model (Fig. 2a). Because P_i_ release from the F-actin core is a slow, stochastic event with a half-time τ >100 seconds, it is unattainable to study the P_i_ escape path using conventional MD simulations. Instead, we developed an enhanced-sampling simulation protocol based on meta-dynamics, which applies a history-dependent repulsive potential on the P_i_ Cartesian coordinates to progressively drive it out of the nucleotide-binding site^32^, without favoring any egress route *a priori* (see Methods for details).

**Fig. 2.**
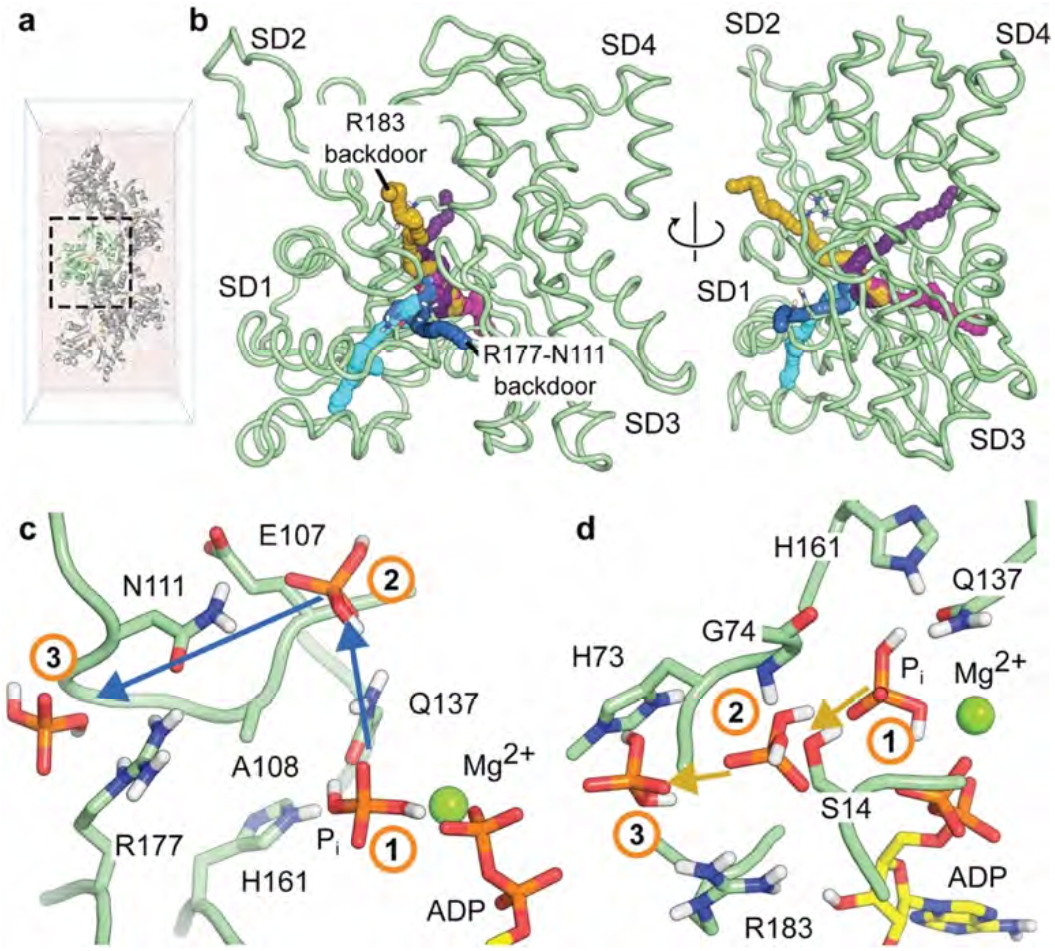
Enhanced sampling simulations reveal several P_i_ release paths from the F-actin core. **a** Simulation box containing an explicitly solvated actin pentamer. The core actin subunit is shown in pale green. **b** Typical P_i_ egress pathways from actin core obtained by enhanced sampling. Actin subdomains are annotated. The two plausible egress paths that were analyzed further are annotated. Implausible P_i_ release pathways are further shown in Supplementary Fig. 4. **c, d** Close-up views of plausible P_i_ egress paths. Time-ordered representative P_i_ positions from enhanced sampling trajectories are shown and connected by arrows indicating the direction of P_i_ movement. Close-up views are depicted for the potential R177-N111 backdoor path (panel **c)** and for on the potential R183 backdoor path (panel **d)**.

Using this approach, we collected dozens of P_i_ release events from the F-actin core and identified sterically accessible egress pathways (Fig. 2b). We then analyzed the predicted pathways for physical plausibility and first dismissed all pathways that entailed unrealistic distortions of F-actin or the nucleotide (Supplementary Fig. 4c-e). Secondly, we made the assumption that P_i_ exits the F-actin interior near the binding sites of phalloidin and jasplakinolide, as both toxins strongly inhibit P_i_ release^33–35^. This analysis resulted in two remaining egress pathways that are physically plausible. In the first one, a side-chain movement of Q137 allows P_i_ to move into a hydrophilic pocket between E107 and H161, where it also interacts with the Pro-rich loop residues. From there, P_i_ is observed to escape either by disrupting the R177-N111 hydrogen bond (leading to an open backdoor, similar to the conformation adopted by the barbed end) (Fig. 2c, Supplementary Video 2), by leaving close to residues N115 and R116, or by exiting near residues T120 and V370. Of note, phalloidin and jasplakinolide stabilize the R177-N111 interaction (Supplementary Fig. 2c), but would not interfere with P_i_ egress near N115-R116 or T120-V370, suggesting that the exit path that requires the disruption of the R177-N111 interaction is the most probable. In the second plausible pathway, P_i_ first breaks the hydrogen bond between S14 and G74 to enter a pocket between H73 and R183. Then, P_i_ escapes to the intra-filament space upon breaking the strong electrostatic interaction with R183 (Fig. 2d). Interestingly, the disruption of the S14-G74 hydrogen bond was previously proposed to play a role in P_i_ release, albeit *via* a different mechanism^31^. Furthermore, phalloidin and jasplakinolide would prevent the opening of the H73-R183 pocket. Thus, in addition to the predicted route, there exists another physically realistic egress pathway for P_i_.

### Actin filaments harboring the N111S mutation release P_i_ rapidly

To experimentally probe the two possible P_i_ release pathways from the F-actin core, we mutated key residues in β-actin that pose a barrier for P_i_ release in each pathway. For the first pathway, we introduced the N111S mutation to potentially disrupt the hydrogen-bonding network of the R177-N111 backdoor. For the second pathway, we aimed to destabilize the R183-mediated backdoor by R183W and R183G mutations. Importantly, all actin mutants studied here are associated with human diseases; the R183W mutation was identified in β-actin of patients suffering from deafness, juvenile-onset dystonia and development malfunctions^36, 37^, whereas the R183G and N111S mutations have been found in α-actin of nemaline myopathy patients^38, 39^, highlighting the relevance of these actin variants.

We developed a fluorescence-based assay to synchronously monitor seeded actin polymerization using pyrene fluorescence and subsequent P_i_ release via a fluorescent phosphate sensor in the same experiment (Fig. 3a). This allowed us to determine the respective reaction rates by fitting the data to a kinetic model (see Methods for details). Because we performed the experiments at actin concentrations (10 μM) that allow for rapid filament growth and seeded the reaction with spectrin-actin seeds to circumvent slow nucleation, we effectively monitored P_i_ release from internal F-actin subunits and not from the barbed end. The assay revealed that wild type β-actin released P_i_ at a slow rate k_-Pi_ of 0.0065 s^-1^ (half time τ ∼107 s) (Fig. 3b), which falls within the range of values previously reported for rabbit skeletal α-actin^10, 20, 24^, indicating that slow P_i_ release after polymerization and ATP-hydrolysis is a feature conserved between mammalian actin isoforms. R183G- and R183W-actin released P_i_ at a slightly increased rate of, respectively, 0.0117 s^-1^ and 0.0190 s^-1^, corresponding to a 1.8- and 2.9-fold increase compared to wild-type β-actin (Fig. 3b). Strikingly, N111S-actin exhibited ultrafast P_i_ release kinetics without appreciable delay after polymerization. In fact, the reaction time-courses of P_i_ release slightly outpaced the observed polymerization kinetics (Fig. 3b) making it impossible to determine an exact P_i_ release rate for N111S-actin. However, when estimated conservatively (see Methods, Supplementary Fig. 5c, d), N111S-actin releases P_i_ at a rate ≥0.1 s^-1^, which is at least 15-fold faster than wild-type actin. Thus, the R183 mutants release P_i_ somewhat faster but still display the characteristic delay between polymerization/ATP-hydrolysis and P_i_ release, whereas in contrast, N111S-actin appears to release P_i_ without appreciable delay.

**Fig. 3:**
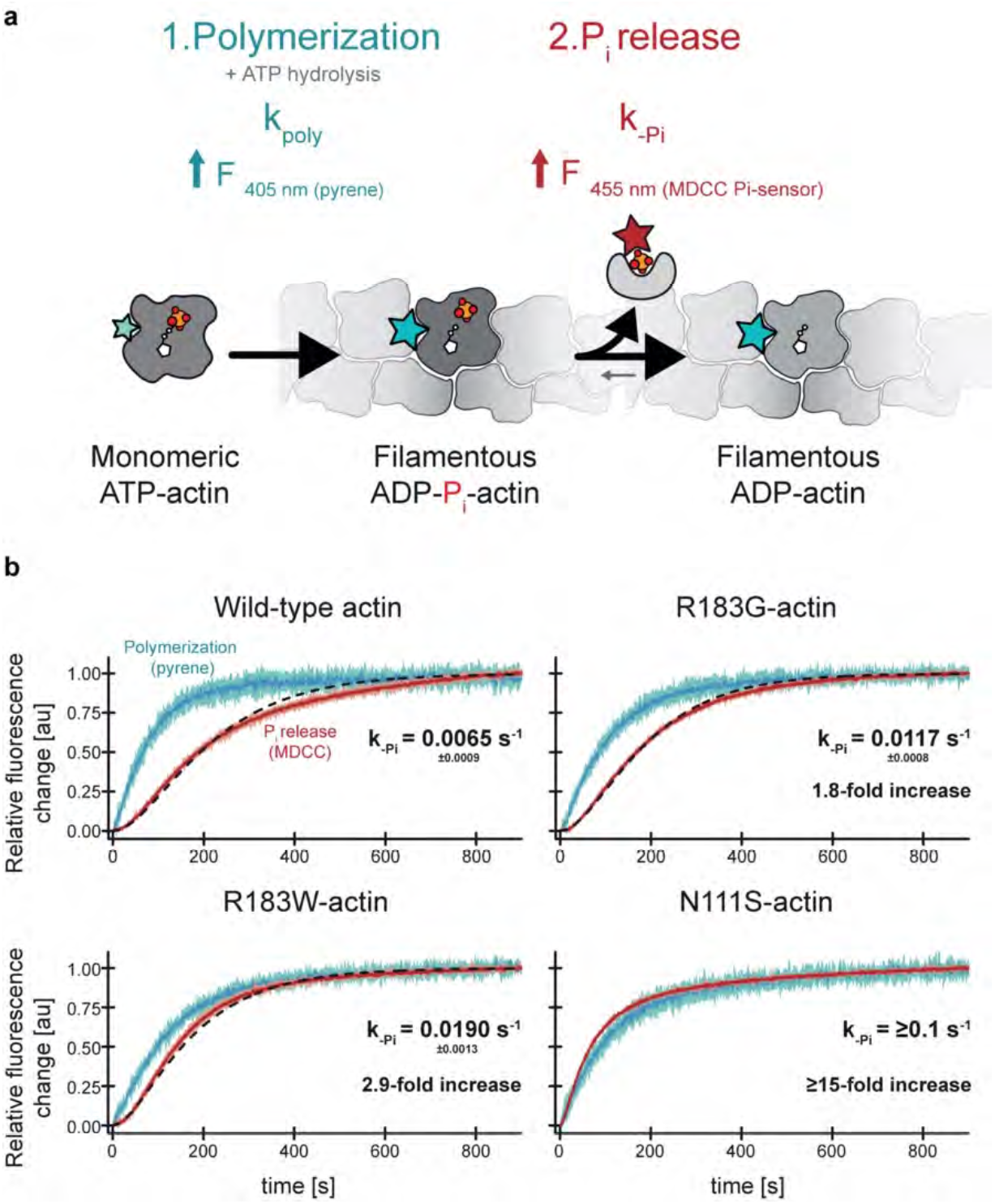
Biochemical characterization of P_i_ release from actin filaments. **a** Scheme of the synchronous measurement of actin polymerization and subsequent P_i_ release as measured by the fluorescence intensity increase of pyrene (cyan) and MDCC-PBP (red), respectively. **b** Time courses of the normalized fluorescence intensities from 10 µM actin (either wild-type or mutants as indicated) containing 1.5% wild-type, pyrene α-actin (cyan) and 30 µM MDCC-PBP (red) seeded with 160 nM spectrin-actin seeds after initiation of polymerization (t=0s). Dark colors indicate averages from three independent experiments whereas light color areas indicate SD. The black dashed lines correspond to fits of the phosphate release data to a kinetic model (see Methods). Rates of P_i_ release and relative rate enhancement over wild-type actin as determined either from fits to a kinetic model (wild-type, R183G and R183W actin) or estimated from kinetic simulations (N111S, see Methods, Supplementary Fig. 5c, d) are depicted in each graph.

### Structural basis for the ultrafast P_i_-release kinetics of N111S-actin

To structurally understand the differences in P_i_ release rates between the R183W and N111S mutants, we determined the filament-core structures of these variants in the Mg^2+^-ADP-bound state at ∼2.3-Å by cryo-EM (Fig. 4a, b, Supplementary Table 1, Supplementary Figs 6, 7, 8a, b). We could model hundreds of water molecules bound to the actin filament in both structures, as well as the exact rotameric positions of many amino-acid sidechains (Supplementary Fig 8a-c). Globally, we observed no major differences between the two β-actin mutants and Mg^2+^- ADP-bound α-actin filaments (pdb 8a2t, rmsd <0.7 Å) (Supplementary Fig. 8d) but, importantly, we identified small but impactful rearrangements.

**Fig. 4.**
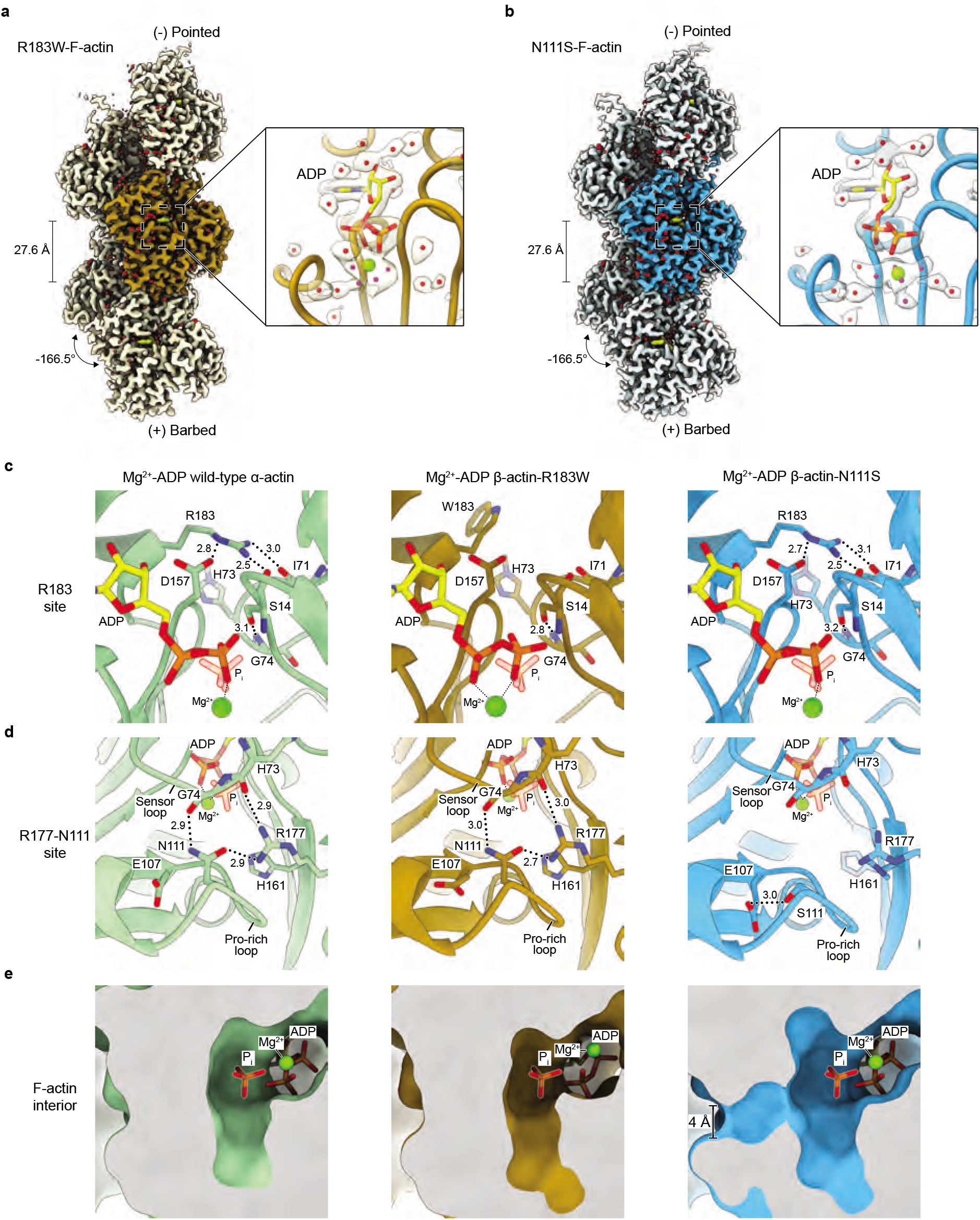
High-resolution cryo-EM structures of R183W- and N111S-F-actin. **a, b** Sharpened cryo-EM density maps of R183W-F-actin **(a)** and N111S-F-actin **(b)** in the Mg^2+^-ADP nucleotide state. Five filament subunits are shown. For the R183 mutant, the central subunit is colored gold, whereas the central subunit of N111S-F-actin is colored blue. The helical rise and twist are annotated. Densities corresponding to water molecules are red. For each mutant, a zoom of the nucleotide-binding pocket with corresponding cryo-EM densities for the nucleotide (depicted yellow), the Mg^2+^-ion (green) and water molecules (red) is shown. Water molecules that directly coordinate the Mg^2+^-ion are colored magenta. **c-e** Structural comparison between filamentous wild-type α-actin (pdb 8a2t, left panel, in green), R183W-β-actin (middle panel, in gold) and N111S- β-actin (right panel, in blue) in the Mg^2+^-ADP state. The panels depict the amino acid environment near the location of residue-183 **(c)**, the amino-acid environment near residues 177 and 111 **(d)**, and a slice through the F-actin interior near the P_i_ binding site **(e)**. In panels **c** and **d**, F-actin is shown as cartoon and amino acids are shown as sticks and are annotated. In panel **e**, F-actin is depicted in surface representation. Water molecules are omitted from panels **c – e**. Although the depicted structures adopt the ADP state, P_i_ from pdb 8a2s (F-actin in the Mg^2+^-ADP-P_i_ state) is fitted and shown semi-transparently to emphasize the P_i_-binding site.

We first examined the atomic arrangement near the mutated residues. In wild-type F-actin structures, the R183 sidechain interacts with D157 and the carboxyl moieties of S14 and I71 (Fig. 4c). As expected for R183W-F-actin, these interactions are abolished and the W183 sidechain points away from the mentioned residues (Fig. 4c). Nevertheless, the S14-G74 hydrogen bond remains intact and we did not observe an open cavity near W183 through which P_i_ could escape (Fig. 4c); the R177-N111 backdoor is also intact in the R183W-F-actin structure (Fig. 4d, e).

In N111S-F-actin, the introduced mutation induces a more drastic conformational change; residue S111 is too short to interact with R177 and G74. Instead, S111 hydrogen bonds with E107, the backbone of A108 and a water molecule within its local environment in the Pro-rich loop (Fig. 4d, Supplementary Fig. 8c). Additionally, the density for the sidechain of R177 is fragmented (Supplementary Fig. 8c), indicating that the residue is more flexible than in wild-type F-actin structures. As a result, the interaction between the sensor loop and Pro-rich loop is disrupted and a ∼4-Å diameter hole is observed, yielding an open backdoor (Fig. 4d, e). Hence, the conformation of F-actin subunits harboring the N111S mutation is reminiscent of the arrangement of the ultimate subunit of the barbed end structure (Fig. 1d). Further consistent with the barbed end structure, we identified that H161 adopts a mixture of rotameric states and that it partially adopts a G-actin-like conformation (Fig. 4d, Supplementary Fig. 8c). This observation suggests that repositioning of the H161 sidechain represents a key feature of backdoor opening. Conversely, the S14-G74 interaction is not affected in N111S-F-actin (Fig. 4c), indicating that disruption of this interaction is not required to open the backdoor as previously proposed^31^. Thus, although the backbone atom shifts in N111S-actin are minor when compared to wild-type actin (<1 Å), sidechain rearrangements result in a broken hydrogen-bonding network and an open backdoor (Fig. 4d, e, Supplementary Fig. 8c, Supplementary Video 3), defining the structural basis for the ultrafast P_i_ release kinetics of N111S-F-actin.

We furthermore inspected the conformation of ADP and the associated Mg^2+^ ion in both mutant structures. In the N111S-F-actin structure, the nucleotide arrangement is essentially the same as that in Mg^2+^-ADP wild-type α-actin; the Mg^2+^ ion resides beneath the β-phosphate of ADP and furthermore interacts with five water molecules (Fig. 4b, Supplementary Fig. 8e, Supplementary Video 3). Interestingly, in R183W-F-actin, the Mg^2+^ changes position so that it is directly coordinated by both α- and β-phosphates, as well as by four water molecules (Fig. 4a, Supplementary Fig. 8e). Although residue R183 does not directly interact with ADP in wild-type F-actin, it resides in close proximity to the nucleotide and, importantly, it is positioned in a negatively-charged cluster of acidic residues (Supplementary Fig. 9). Hence, R183W-F-actin harbors a more negatively charged nucleotide-binding site (Supplementary Fig. 9), which may explain why the positively charged Mg^2+^ repositions in the mutant structure to compensate the charge imbalance. Thus, R183W-F-actin displays an altered nucleotide-binding site but does not reveal any conformational changes that would allow for P_i_ egress, indicating that the R183-backdoor does not encompass the dominant P_i_ escape path in wild-type actin. Although we cannot exclude that this release path is marginally sampled, we propose that, alternatively, P_i_ still mainly exits through the R177-N111 backdoor in the R183 mutants. In that scenario, the more negatively charged nucleotide-binding pocket may explain why the negatively charged P_i_ is released slightly faster from R183W- and R183G-actin compared to wild-type actin (Fig. 3b, c).

### N111S-actin filaments display dramatically shortened ADP-P_i_ caps

Next, we hypothesized that N111S-actin, which releases P_i_ rapidly in bulk assays and adopts an open backdoor, should also display strong differences in nucleotide state distribution at the single filament level compared to wild-type actin. Specifically, we reasoned that filaments harboring the N111S mutation should form a drastically shortened ADP-P_i_ cap, which we tested in microfluidic flow-out assays using total internal reflection fluorescence (TIRF) microscopy (Fig. 5a, Supplementary Video 4). Filaments were elongated from surface-immobilized spectrin actin seeds using either wild-type- or N111S-actin and then rapidly switched to depolymerization with buffer lacking soluble actin (see Methods). For wild-type β-actin, the speed of filament depolymerization after flow-out gradually increased over a few minutes to converge to a maximal rate (Fig. 5b, c). Since ADP-P_i_-bound actin depolymerizes at slower rates than ADP-bound actin, this change is caused by the filament depolymerizing region slowly maturing from an ADP-P_i_-rich to an ADP-rich composition through P_i_ release. The measured depolymerization velocities are well described by a kinetic model (Fig. 5d, see Methods) with parameters similar to those previously measured from single-filament assays on skeletal α-actin^20^, as well as from our own bulk measurements (Fig. 3b, Fig. 5d). Strikingly, we observed that filaments grown from N111S actin depolymerized at a high velocity (v_depol,ADP_ = 14.80 s^-1^) (Fig. 5b, c). More importantly, we found no appreciable change in the depolymerization velocity after buffer flow-out (Fig. 5c), indicating that the ADP-P_i_ to ADP- actin transition is very fast and not captured within the resolution of our experiment. This allowed us to estimate a lower bound for the rate of phosphate release from the filament interior k_-Pi_ ≥ 0.154 s^-1^ for N111S actin (Fig. 5d, Supplementary Fig. 5e). Hence, our data reveal in high mechanistic detail that the rapid rate of P_i_ release indeed results in drastically shortened ADP-P_i_ caps in N111S-actin filaments.

**Fig. 5.**
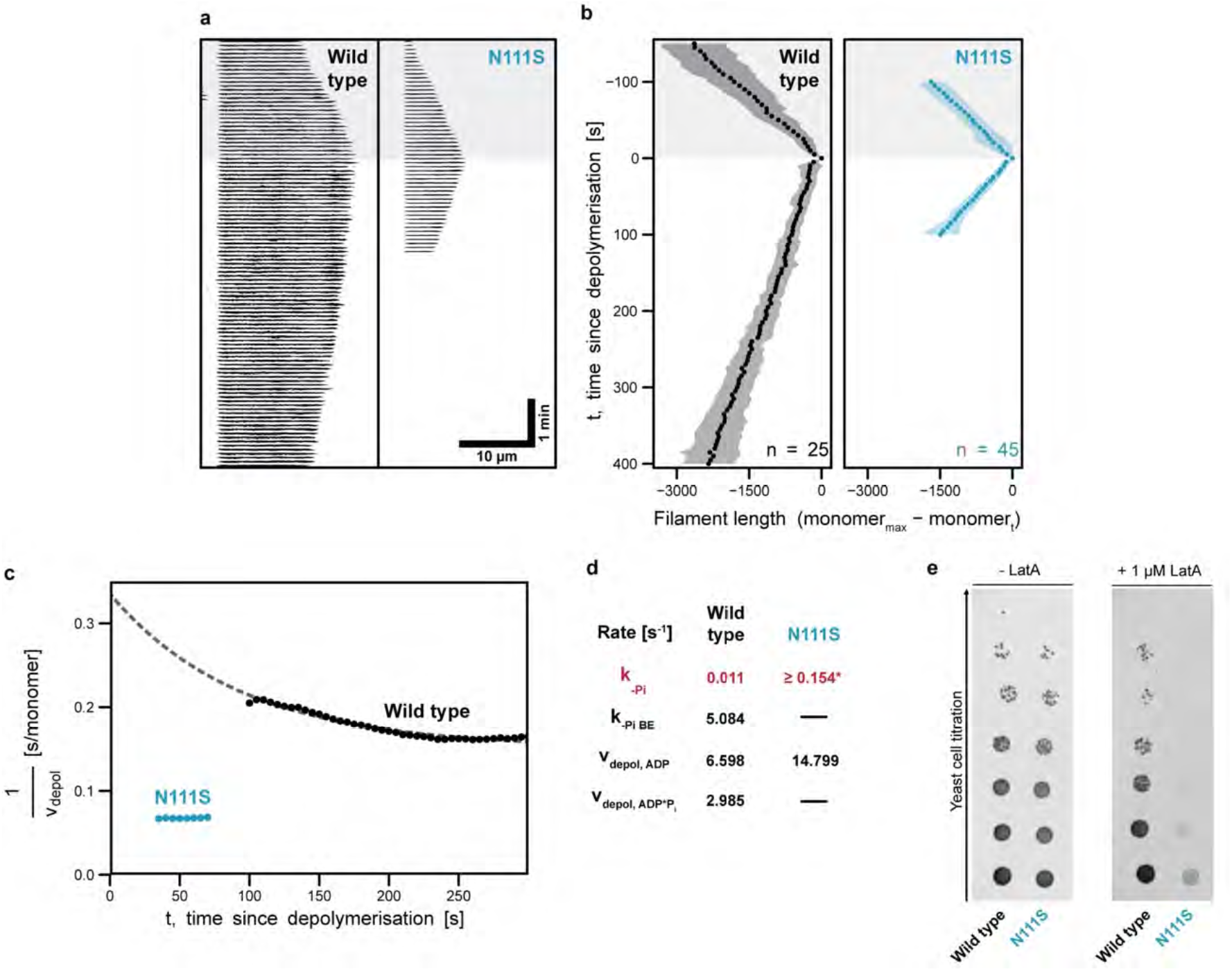
Effects of the N111S mutation on single actin filaments *in vitro* and on yeast growth *in vivo*. **a** Timelapse TIRF imaging of single actin filaments (black, visualized by Lifeact- AlexaFluor-488) polymerized from spectrin-coated surfaces in microfluidic flow chambers. The vertical and horizontal axis indicate time and filament length, respectively. Polymerization and depolymerization phases are demarcated by the grey shaded and unshaded areas respectively. **b** Filament lengths tracked over the course of polymerization (grey shaded area) and depolymerization (unshaded area). Points indicate average filament length calculated from 25 and 45 filaments polymerized from either wild-type- or N111S-actin, respectively. Shaded areas indicate SD. Wild-type and N111S filaments were analyzed from 6 and 12 independent experiment, respectively. **c** The inverse of the instantaneous depolymerization velocity (1/v_depol_) calculated from the average filament length in e as a function of time after depolymerization. The dashed line indicates a fit to a kinetic model (see Methods). **d** Rate constants of phosphate release from the filament interior and barbed end (k_-Pi_ and k_-Pi BE_) and depolymerization velocities of ADP-P_i_ or ADP subunits from the filament end (v_depol,ADP_ and v_depol,ADP-Pi_) determined from fits obtained in c. * indicates lower bound estimate for N111S (see Methods, Supplementary Fig. 5e). **e** Growth phenotype assay with yeast expressing either wild-type or N111S variants of *S. cerevisiae* actin. The two yeast strains were grown in the absence and presence of 1 μM Latrunculin A (LatA).

To also investigate the effect of the N111S mutation *in vivo*, we compared the growth rates of yeast strains expressing either wild type or N111S variants of *S. cerevisiae* actin. Under normal conditions, we observed no major differences in growth phenotype between the two strains (Fig. 5e). However, when exposed to the toxin Latrunculin A, which sequesters G-actin and accelerates F-actin depolymerization^40^, the yeast strain expressing N111S-actin displayed a dramatically reduced growth rate compared to that of the strain expressing wild-type-actin (Fig. 5e). This high sensitivity to Latrunculin A-induced stress suggests that N111S-actin filaments are more labile and prone to depolymerization. Thus, the phenotype observed for *S. cerevisiae* N111S-actin *in vivo* is in line with our *in vitro* experiments on human-β N111S- actin, where we observed faster P_i_ release and filament depolymerization rates compared to wild-type actin.

Taken together, our experiments provide strong evidence that the first P_i_ egress pathway identified by enhanced sampling MD, encompassing the R177-N111 backdoor, is the dominant route for the exit of P_i_ from the F-actin interior. Hence, P_i_ is released from the core and barbed end of F-actin through similar exit routes, although F-actin subunits in the filament core require additional conformational rearrangements to transiently open the R177-N111 backdoor.

### Visualization of the transient state that allows for P_i_-release from the F-actin core

Finally, we sought to understand how P_i_ is released through the R177-N111 backdoor from the wild-type F-actin core. Our enhanced-sampling MD suggested P_i_ escape pathways through the backdoor (Fig. 2c), but this protocol is aggressive and may not reflect a realistic order of molecular motion. Therefore, we first ran unbiased MD simulations to investigate the backdoor conformation in F-actin. The simulations revealed the disruption and reformation of the R177- N111 interaction several times over a 1.1 μs period (Supplementary Fig. 10a). However, this never resulted in a functionally open backdoor (Supplementary Fig. 10b), indicating that more rearrangements are required for P_i_ to escape, which were not captured in the timeframe of our unbiased simulations. This is in line with the very slow P_i_ release kinetics of wild-type actin (Fig. 3b).

Therefore, to gain further mechanistic insights into the formation of this transient state of the F-actin core that allows for P_i_ release, we turned to steered-MD (SMD) simulations. We performed 40 SMD simulations of 100 ns each, in which selected regions of the backbone of an F-actin core protomer (with either charged or neutral meH73) were transitioned to the barbed end conformation, which is a prototypical example of a functionally open backdoor (see Methods). The majority of simulations showed a Pro-rich loop movement and the disruption of the R177-N111 hydrogen bond within 25 ns (Fig. 6a, b, Supplementary Fig. 11, 12), representing the first major step of backdoor opening. The rotation of H161 to a G-actin like conformation was also observed in many simulations and generally occurred after disruption of the R177-N111 interaction within 40-50 ns (Fig. 6b, c, Supplementary Fig. 11, 12), while inter-subunit contacts within the filament remained intact. We then evaluated the P_i_ release efficiency of structures extracted along these key timepoints in the SMD trajectories using our previously introduced metadynamics-based protocol. While we did not observe a major effect of the charge of meHis73 on backdoor opening, our simulations revealed a high probability (p_BD_) (Fig. 6c, Supplementary Fig. 12a, b, 13, see Methods) for P_i_ to escape through the backdoor when the R177-N111 interaction was disrupted and the H161 sidechain was flipped, suggesting that both events are required to stabilize an open backdoor conformation. Thus, although only the backbone movement was steered in our SMD setup, the simulations elucidate the sidechain rearrangements that lead to the transient opening of the backdoor in the F-actin core (Fig. 6).

**Fig. 6.**
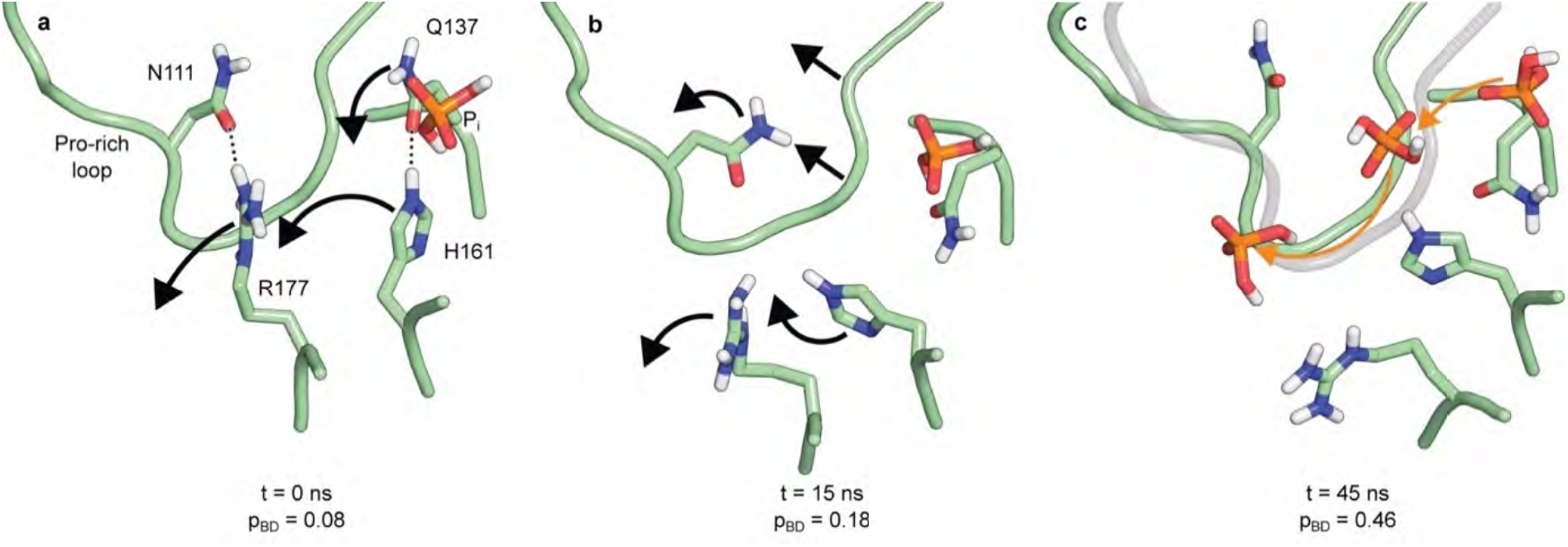
Backdoor opening mechanism revealed by SMD simulations. **a-c** Selected frames from the SMD trajectory. Dotted lines indicate hydrogen bonds. Arrows indicate side chain and backbone movements involved in the opening of the backdoor. Time *t* and P_i_ release propensity p_BD_ along the trajectory are shown. In panel **c,** the orange arrows indicates the direction of P_i_ movement through the open R177-N111 backdoor observed in a representative enhanced sampling simulation.

## Discussion

By combining cryo-EM with *in vitro* reconstitution and MD simulations, we have elucidated how P_i_ is released from F-actin (Fig. 6, 7) and that the hydrogen bonding network between the Pro-rich loop, sensor loop and R177-strand forms the predominant backdoor. At the barbed end, the door is open for P_i_ release in the ultimate subunit, indicating that P_i_ can egress without large protein rearrangements. However, the backdoor is closed in F-actin core subunits and only opens transiently through a high-energy state. The opening of the door is likely initiated by a stochastic disruption of the hydrogen bonding network and further induced by the change of rotameric position of H161 to a G-actin-like conformation. After P_i_ egress, H161 flips back to its original position and the hydrogen bonding network is re-established, allowing F-actin to adopt its low-energy state with a closed backdoor. In the N111S-actin mutant, which is linked to nemaline myopathy, the backdoor is predominantly open in all subunits because the introduced S111 sidechain is too short to maintain the interactions that keep the door closed. Hence, N111S-F-actin releases P_i_ rapidly upon polymerization, thereby drastically reducing the fraction of ADP-P_i_-bound subunits in the filament. Thus, our data provide conclusive evidence that the P_i_ release rate from F-actin is controlled by steric hindrance through the backdoor rather than by the disruption of the ionic bond between P_i_ and Mg^2+^ at the nucleotide-binding site as previously proposed^27^.

**Fig. 7.**
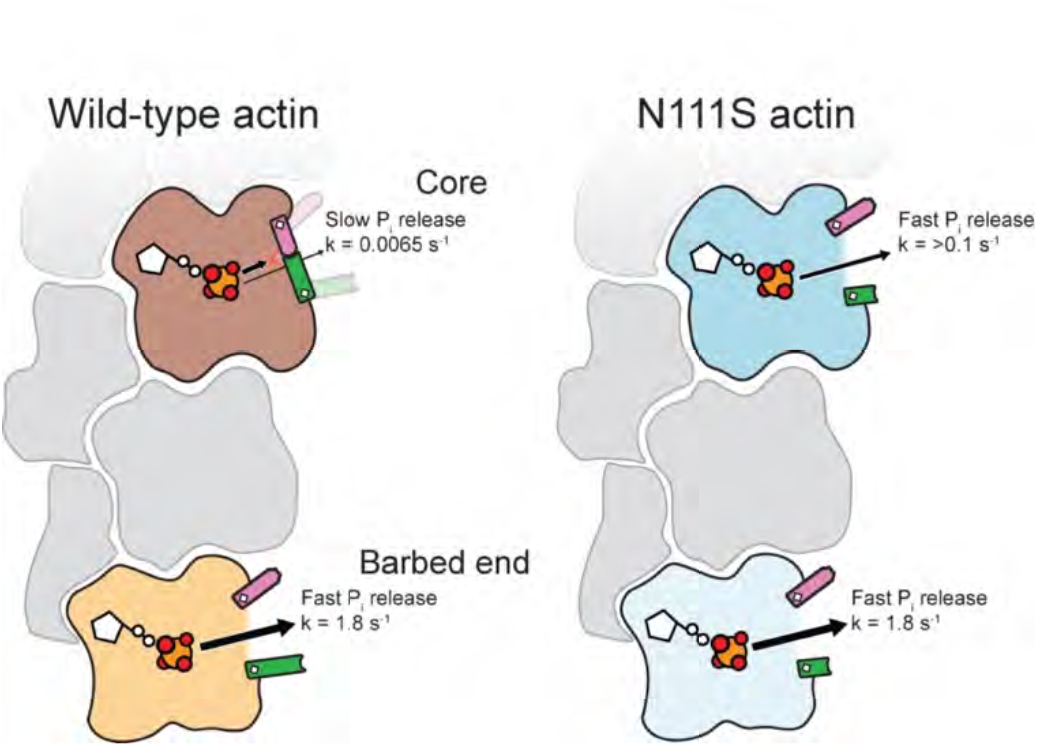
Cartoon model of P_i_ release from wild-type and N111S-F-actin. In wild-type F-actin, subunits that reside in the filament core predominantly adopt a closed backdoor and hence, release P_i_ at slow rates. In the ultimate subunit at the barbed end, the backdoor is open, leading to a 300-fold faster P_i_ release during actin depolymerization. In N111S-F-actin, the amino-acid substitution results in an open backdoor in internal subunits, leading to increased P_i_ release rates.

The open backdoor arrangement in the ultimate subunit at the barbed end explains how remaining P_i_ can be rapidly released during filament depolymerization. In contrast, under filament growth conditions, the barbed end growth velocity (∼10 – 500 monomers s^-1^, dependent on the actin concentration)^41^ is much faster than the ATP hydrolysis rate of F-actin (0.3 s^-1^)^9^, indicating that barbed end subunits effectively only adopt the ATP state before becoming internal subunits. During the G- to F-actin transition, the sidechain of residue H161 flips towards ATP, which triggers the relocation of water molecules near the nucleotide and, as a result, creates a favorable environment for ATP-hydrolysis^7, 8^. In contrast, in our barbed end structure, H161 adopts its G-actin like rotameric position and points away from the nucleotide in the ultimate subunit, which suggests that, regardless of the filament growth velocity, the last subunit only becomes ATP-hydrolysis competent when another actin subunit is added to the filament. This provides implications that the complete G- to F-form transition of a given actin subunit should be considered as a multi-step process, which not only encompasses the initial incorporation of that subunit into the filament, but also requires the subsequent binding of the next actin subunit. Formal proof for such a mechanism will require future structural investigations of the barbed end of F-actin in the ATP state.

Can our results explain how the N111S mutation leads to nemaline myopathy, a disease affecting skeletal muscle α-actin? Although actin filaments in striated muscle are expected to undergo less turnover than cytoplasmic actin isoforms, it is well established that actin-severing proteins such as cofilins are expressed in muscle sarcomeres to control actin (thin) filament length during sarcomerogenesis and actin turnover^42–44^. Since cofilins preferably sever ADP- over ADP-P_i_-bound F-actin, we propose that the ultrafast P_i_-release kinetics of N111S-actin may contribute to the pathophysiology of the disease in these patients. Moreover, it will be interesting to study the effects of the N111S mutation in actin in non-muscle tissue. Because N111S actin does not majorly populate the ADP-P_i_ state, it could represent a unique tool for investigating the role of this metastable actin state both *in vivo* and *in vitro*. In general, we envision that our approach of studying actin mutants R183W and N111S, where we combined biochemical experiments with high-resolution cryo-EM, will also be instrumental in elucidating the molecular mechanisms of other disease-associated actin mutants. Specifically, visualizing the mutations’ impact on the F-actin atomic structure will allow for the formulation of new hypotheses on how the mutations affect cellular processes and how this is linked to disease. Therefore, this approach may ultimately contribute to the development of new therapeutic strategies for the treatment of human diseases that are characterized by mutations in actin genes.

## Methods

### DNA constructs and yeast strains

Throughout the manuscript, we used human β-actin amino-acid numbering that is consistent with the numbering in the corresponding UniProt entry (P60709, ACTB_HUMAN), with the initiator methionine numbered as residue 1, even though this methionine is cleaved off during actin maturation. Hence, the used amino-acid numbers for β-actin are in register with the sequences of mature human and rabbit skeletal α-actin, as well as *S. cerevisiae* actin, facilitating a direct comparison between all actin isoforms used in this study.

The plasmid for the expression of recombinant, human β-actin (p2336 pFL_ACTB_C272A) was described previously^45^. All mutations were introduced *via* QuikChange PCR, using p2336 as template. All β-actin constructs contain the C272A substitution (including the protein referred to as wild type), as C272 is prone to oxidation in aqueous solutions^46^. The equivalent residue in human skeletal α-actin is also alanine. The FH2 domain of *M. musculus* mDia1 (mDia1_FH2_) (amino acids 750-1163) was cloned into an pETMSumoH10 expression vector by Gibson assembly.

*S. cerevisiae* strains were grown in minimal medium containing yeast nitrogen base without amino acids (Difco) containing glucose and supplemented with tryptophane, adenine, histidine, and/or uracil if required. *S. cerevisiae* strains were transformed using the lithium-acetate method^47^. Yeast actin mutagenesis was performed as described previously^48^. Yeast viability in the presence or absence of Latrunculin A was analyzed by drop test assays: 5-fold serial dilutions of cell suspensions were prepared from overnight agar cultures by normalizing OD_600_ measurements, then plated onto agar plates and incubated at 30°C for 2 days.

### Protein expression and purification

Native bovine, cytoplasmic β-actin and γ-actin mixture was purified from bovine thymus tissue as described previously^41, 49^. Bovine β and γ-actin both display 100% amino-acid sequence identity to their corresponding human orthologs.

Human cytoplasmic β-actin variants were recombinantly expressed as fusion proteins, with thymosin β4 and a deca-His-tag fused to the actin C-terminus^45, 50^. β-actin was expressed in BTI-Tnao38 insect cells using baculovirus infection. ∼48 hours after infection, the cells were pelleted (5000*g*, 10 min) and stored at −80 °C until use. On the day of purification, the insect cells were thawed, resuspended in Lysis Buffer containing 10 mM Tris pH 8, 50 mM KCl, 5 mM CaCl_2_, 1 mM ATP, 0.5 mM TCEP and cOmplete protease inhibitor (Roche), and subsequently lysed using a fluidizer. The lysed cells were subjected to ultracentrifugation (100,000*g*, 25 min) to remove cell debris. The supernatant was filtered and then loaded onto a 5 ml HisTrap FF crude column (Thermo). The column was washed with 10 column volumes of lysis buffer supplemented with 20 mM imidazole. The fusion protein was eluted from the column using a linear imidazole gradient (20 mM to 160 mM in 10 min). The eluted fusion-protein was then dialyzed overnight in G-buffer (5 mM Tris pH 8, 0.2 mM CaCl_2_, 0.1 mM NaN_3_, 0.5 mM ATP, 0.5 mM TCEP), followed by incubation with chymotrypsin (w/w) for 20 min at 25 °C to cleave thymosin β4 and the deca-His-tag from β-actin. Proteolysis was terminated by the addition of 0.2 mM PMSF (final concentration) and the mixture was applied to a clean HisTrap FF crude column. The flowthrough was collected and β-actin was polymerized overnight by the addition of 2 mM MgCl_2_ and 100 mM KCl (final concentrations). The next morning, actin filaments were pelleted by ultracentrifugation (210,000*g*, 2hrs) and resuspended in G-buffer. β-actin was depolymerized by dialysis in G-buffer for 2 – 3 days. Afterwards, the protein was ultracentrifuged (210,000*g*, 2hrs) to remove any remaining filaments. The supernatant was, if necessary, concentrated in a 10 kDa concentrator (Amicon) to 2 – 3 mg/ml, flash frozen in liquid nitrogen in 50 µl aliquots and stored at −80 °C until further use.

Native, rabbit 𝘢-actin was purified from muscle acetone powder and labelled with pyrene (N-1-Pyrene-Iodoacetamide). 10 g of rabbit muscle acetone powder was mixed with 200 mL G-Buffer and stirred for 30 min at 4 °C. Later, the solution was centrifuged at 10,000 rpm in a GSA rotor for 1 hr at 4 °C to separate the supernatant containing actin from the debris. Actin polymerization was initiated in the supernatant by adding Polymerization Buffer (50 mM KCl, 1.5 mM MgCl_2_, 1 mM EGTA, 10 mM imidazole pH 7) and an additional 0.1 mM ATP. The solution was allowed to polymerize at room temperature for 1 hr while being stirred, after which the solution was moved to 4 °C to polymerize for another hour. Later, the concentration of KCl in the solution was increased to 800 mM and stirred for another 30 min. After a total of 2.5 hr of polymerization the solution was centrifuged at 35,000 rpm in a Ti45 rotor for 1 hr at 4 °C and the actin pellet was resuspended in F-Buffer (2 mM Tris pH 8.0, 50 mM KCl, 1.5 mM MgCl_2_, 1 mM EGTA, 10 mM imidazole, 0.1 mM CaCl_2_, 0.2 mM ATP, 0.01% NaN_3_, 0.5 mM TCEP).

Polymerized actin was reduced by adding 1 mM TCEP and centrifuged at 80,000 rpm in a TLA-110 rotor for 30 min at 4 °C. The resulting actin pellet was resuspended in labelling buffer (50 mM KCl, 1.5 mM MgCl_2_, 1 mM EGTA, 10 mM imidazole pH 7, 0.2 mM ATP) to which pyrene was added in 5 times molar excess of actin and incubated in ice for 2 hr. Reaction was quenched by adding 10 mM DTT and dialyzed in G-Buffer to depolymerize actin.

For the purification of heterodimeric DNase-I:actin complexes, deoxyribonucleaseI from bovine pancreas (DNase I, Serva, Cat.# 18535.02) was dissolved at a concentration of 666 μM (20mg/ml) in 1xKMEI Buffer (0.5 mM ATP, 1 mM TCEP, 50 mM KCl, 1.5 mM MgCl_2_, 1 mM EGTA, 10 mM imidazole pH 7) containing 1x complete protease inhibitors and 1mM PMSF. 4ml of 90uM filamentous β,γ-actin were mixed with 1.1ml of 666 μM DNase I, resulting a 1:2 molar ratio, and depolymerized by dialyzing for one week against G-buffer. The dialyzed sample was centrifuged for 30 min at 80.000 rpm in a TLA110 rotor and the supernatant was gel filtered into G-Buffer over a Superdex 200 16/600 column (Cytiva). Fractions corresponding to the heterodimeric DNase-I:actin complex were pooled, concentrated and stored at 4°C for up to three months.

mDia1_FH2_ was expressed with an N-terminal 10xhis-SUMO3-tag in *E. coli* BL21 Star pRARE cells for 16 hr at 18°C. The cells were lysed in Lysis Buffer-v2 (50 mM NaH_2_PO_4_ pH 8.0, 400 mM NaCl, 0.75 mM β-mercaptoethanol, 15 μg/ml benzamidine, 1xcomplete protease inhibitors, 1 mM PMSF, DNaseI) and the protein was purified by IMAC using a 5 ml HisTrap column. The protein was eluted using Elution Buffer (50 mM NaH_2_PO_4_ pH 7.5, 400 mM NaCl, 400 mM imidazole, 0.5 mM β-mercaptoethanol) in a gradient and the 10xhis-Sumo3-tag was directly cleaved using SenP2 protease overnight. After cleavage, proteins were desalted into lysis buffer and recirculated over a 5 ml HisTrap column followed by gelfiltration over a Superdex 200 16/600 into Storage Buffer (20 mM HEPES pH 7.5, 200 mM NaCl, 0.5 mM TCEP, 20% glycerol), concentrated, flash frozen in liquid nitrogen and stored at −80°C.

Spectrin-actin seeds were purified and biotinylated as described previously^51^.

### Synchronous measurement of actin polymerization and P_i_ release in bulk assays

On the day of the assay, aliquots of all purified β-actin variants (frozen as G-actin) were thawed and centrifuged at 100,000*g* for 20 – 30 minutes to remove aggregates. To ensure that all variants were in exactly the same buffer, we exchanged the buffer to G-buffer-v2 (5 mM Tris pH 8, 0.2 mM CaCl_2_, 0.1 mM NaN_3_, 0.1 mM ATP, 0.5 mM TCEP) using Micro Bio-Spin® Chromatography Columns (Bio-Rad). For each measurement, a 40 μl G-actin solution containing 30.5 μM unlabeled β-actin variant, 0.5 μM pyrene-labeled, wild-type α-actin and 15 μM P_i_-sensor (MDCC-labeled phosphate binding protein, Thermo Fisher Scientific) was prepared. We confirmed that the presence of trace amounts (1.5%) of pyrene-labeled, wild-type α-actin, which releases P_i_ with slow kinetics, did not significantly contribute to the overall readout by the phosphate sensor, which was dominated by P_i_ release from the actin mutant present in vast excess (98.5%) (Supplementary Fig. 5a). 36 μl of G-actin solution was then mixed with 4 μl 10xME (5 mM EGTA pH 7.5, 1 mM MgCl_2_) and incubated at room temperature for 2 minutes, in order to exchange the ATP-associated divalent cation from Ca^2+^ to Mg^2+^. We then took 36 μl of this solution and mixed it with 64 μl Polymerization buffer (16 mM HEPES pH 7, 160 mM KCl, 3 mM MgCl_2_, 1.5 mM EGTA, 38 μM P_i_-sensor, 160 nM spectrin-actin seeds, 0.1 mg/ml β-casein, 0.1 mM ATP, 0.5 mM TCEP) in a quartz cuvette to start the experiment. This yielded final concentrations of 10 μM actin (1.5% pyrene-labeled), 100 nM spectrin-actin seeds, 100 mM KCl and 30 μM P_i_-sensor. The spectrin-actin seeds were added to ensure rapid polymerization in order to i) minimize potential differences in the polymerization kinetics between the β-actin variants (Supplementary Fig. 5b) and ii) create a pronounced separation between the time courses of polymerization and P_i_ release, at least in the case of wild-type actin. Measurements were carried in a spectrofluorometer (PTI QM-6) under constant excitation at 365 nm and synchronous monitoring at 1s intervals of pyrene (λ = 410 nm) and MDCC (λ = 455 nm) fluorescence intensities, which report on actin polymerization and phosphate release, respectively.

### Determination of P_i_ release rates from bulk assays

Time courses of pyrene or MDCC fluorescence from individual experiments were first normalized by subtracting the minimal signal at the beginning of the experiment and dividing by the maximal signal at saturation at t=900s. Apparent rates of actin polymerization (k_poly_) were then determined by fitting time courses of normalized pyrene fluorescence from individual experiments by a mono-exponential function:

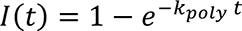

with I(t) being the normalized fluorescence intensity as function of time and k_poly_ being the observed polymerization rate. Average k_poly_ values for each actin variant were calculated from three independent experiments (Supplementary Fig. 5b). To determine rate constants for phosphate release (k_-Pi_), we first averaged time courses of normalized MDCC fluorescence from three individual experiments for each actin variant. This averaged data was fitted by a simple kinetic model using the KinTeK Explorer software (Version 6.3):

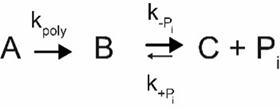

with A monomeric ATP-actin, B filamentous ADP-P_i_-actin, C filamentous ADP-actin. The model contains three kinetic parameters, two of which were fixed. The first-order rate of polymerization (k_poly_), that formally combines the processes of polymerization and ATP hydrolysis was fixed to the experimentally measured polymerization rate for each actin variant (see above). The second-order association rate constant for binding of inorganic phosphate (k_+Pi_) was fixed to 0.000002 µM^-1^ s^-1^ as measured previously for wild-type α-actin and assumed to be the same for all actin variants^11, 25^. This assumption likely does not hold, because the chosen mutations can be anticipated to similarly accelerate both release and binding of P_i_. However, we determined that k_+Pi_ can be varied by more than 1000-fold without significantly affecting the obtained first-order rate of phosphate release (k_-Pi_). More importantly, we can exclude that rebinding of P_i_ contributes significantly under our experimental conditions: P_i_ is i) generated only in minor amounts (10 µM) during the course of the assay and ii) potently sequestered by the phosphate sensor that is present in molar excess (30 µM) and binds P_i_ with 10000-fold higher affinity (K_D_ = 0.1 µM)^52^ compared to actin (K_D_ = 1.5 mM)^11, 25^.

For the N111S mutant, we could not determine the exact rate constant of P_i_ release in this manner, because the average observed rate of P_i_ release slightly exceeded the average observed polymerization rate (Fig. 3b). This should formally not be possible because the latter has to precede the former and the reason for this inversion remains unknown. To nonetheless obtain a conservative estimate for the increase in the P_i_ release rate in this case, we carried out kinetic simulations in KinTek Explorer to systematically explore the dependence of the observed P_i_ release reaction kinetics on the rate enhancement of P_i_ release (Supplementary Fig. 5c, d). This showed that a rate enhancement of P_i_ release by more than 15-fold is required, for the observed P_i_ release rate to fall within the error margin of the observed polymerization rate. Hence, we consider 0.1 s^-1^ the lower bound for the rate of P_i_ release for the N111S mutant.

### Preparation of functionalized glass slides

Functionalized glass slides coated with 5% Biotin-PEG and 95% Hydroxy-PEG were prepared as described^53^. Briefly, high-precision glass coverslips (22 x 60 mm, 1.5H) were asymmetrically cut at one corner using a diamond pen to distinguish the functionalized surface from the non-functionalized one. Glass slides were cleaned by incubating in 3 M NaOH solution for 15 min, rinsed in water and then incubated in freshly prepared Piranha solution (3:2 mixture of 95-97% Sulfuric acid and 30% Hydrogen peroxide) for 30 min. Slides were rinsed with water to remove residual acid and then air dried with nitrogen gas. Dried glass slides were sandwiched with 3 drops of GOPTS (3-Glycidyloxypropyltrimethoxysilane) and then stored in closed Petri dishes, which were further incubated in an oven at 75 °C for 30 min. Sandwiched glass slides were rinsed in acetone and separated with a pair of tweezers. Following separation, the glass slides were rinsed again in fresh acetone. Separated glass slides were air dried with nitrogen gas and placed in pre-warmed Petri dishes with their functionalized surface facing up. The functionalized surfaces were sandwiched with 75 µL of a 150 mg/ml mixture of 95% Hydroxy-amino-PEG (ɑ-Hydroxy-ω-amino PEG-3000) and 5% Biotinyl-amino-PEG (ɑ-Biotinyl-ω-amino PEG-3000) dissolved in acetone. Sandwiched glass surfaces were incubated in closed Petri dishes in an oven at 75 °C for 4 hrs. Following incubation, the glass slides were separated with a pair of tweezers, rinsed multiple times in water and air dried with nitrogen gas.

### Preparation of microfluidic devices

Microfluidic devices were prepared from PDMS (polydimethylsiloxane) casted onto a silicon mold designed to incorporate up to 4 inlets and 1 outlet. A mixture of PDMS and its curing agent (10:1 mass ratio, SYLGARD™ 184 Silicone Elastomer Kit) was thoroughly mixed and poured onto the silicon mold. The PDMS cast was degassed in a desiccator under vacuum for 3 hrs to remove bubbles from the cast. After removal of bubbles, the cast was incubated in an oven at 75 °C for another 4 hrs to complete the curing process. On completion of the curing process, the PDMS microfluidic devices were cut out from the cast, rinsed with isopropanol and air dried with nitrogen gas.

The functionalized glass cover slips and the flow surface of the microfluidic devices were plasma cleaned at 0.35 mbar pressure and 80% ambient air for 3 min. Parts of the functionalized glass cover slips that would eventually line up with the observation chamber on the microfluidic device were protected from plasma treatment by placing blocks of PDMS on the region. The microfluidic device was sealed by placing the PDMS block onto the functionalized glass surface and incubating at 75 °C for 1 hr.

### Microfluidic experiments and TIRF microscopy of single filaments

The surface of a flow channel was prepared by first passivating the surface to avoid unspecific interactions, then coating the biotinylated surface with streptavidin and finally coating the surface with biotinylated spectrin actin seeds. Flow channels were passivated by flowing in 1x KMEI containing 1% Pluronic F-127, 0.1 mg/mL β-casein and 0.1 mg/mL 𝜅-casein. Passivated surfaces were coated with streptavidin by flowing a 1xKMEI containing 0.75 nM streptavidin. Streptavidin coated surfaces were then coated with spectrin actin seeds by flowing in 1x KMEI containing 10 nM biotinylated spectrin actin seeds. Each of the abovementioned steps were interleaved by a wash step by flowing 1xKMEI after each step. All flow steps were carried out at flowrates of 20 µL/min for 5 min each except the streptavidin flow step which was carried out for 2 min. After the surfaces were coated with spectrin seeds, polymerization was started by flowing in assay buffer (0.1 mg/mL β-casein, 1 mM ATP, 1 mM DABCO, 20 mM β-mercaptoethanol, 100 mM KCl, 1.5 mM MgCl_2_, 1 mM EGTA, 20 mM HEPES) containing ∼2.5 µM profilin:actin (2.71 µM wt β-actin or 2.50 µM N111S β-actin) and 50 nM AlexaFluor- 488-Lifeact for visualization of filament growth. The profilin:actin amount was varied to equalize minor differences in the polymerization velocity between the two actin variants. Image acquisition was started immediately after the profilin:actin buffer flow was initiated. Depolymerization was initiated after about 3-5 min of polymerization by stopping the flow of profilin:actin and starting the flow of depolymerization buffer (0.1 mg/mL β-casein, 1 mM ATP, 1 mM DABCO, 20 mM β-mercaptoethanol, 100 mM KCl, 1.5 mM MgCl_2_, 1 mM EGTA, 20 mM HEPES, 50 nM AlexaFluor-488-Lifeact).

Image acquisition was carried out using a TIRF microscope with a 60x objective and a 1.5x zoom lens under TIRF conditions. Time-lapse images were acquired every 5 s with 2% laser power, 2 s exposure time, 150 gain and emitted light was filtered through an emission filter of 525 nm with 50 nm bandpass. All images were acquired at a bit-depth of 16 bits.

### Filament tracking and data analysis

Timelapse TIRF microscopy images of actin filaments were first denoised using the Non-Local Means algorithm implemented with a custom Python script. Lengths of single actin filaments were tracked in time with ImageJ using the JFilament plugin^54^ with the following parameters: alpha = 15, beta = 10, gamma = 20000, weight = 0.5, stretch = 2000, deform iterations = 200, spacing = 1.0, smoothing = 1.01, curve type = open, foreground and background values were assigned by selecting regions of the filament and the surrounding background respectively. Filament lengths tracked from different filaments were averaged and used to calculate depolymerization velocity. Instantaneous depolymerization velocities were calculated by performing local linear regression with a sliding window (size: 20 for wt and 7 for N111S) centered around a time point. An exponential decay function of the form 𝑦 = 𝑎𝑒^!𝑏𝑥^ + 𝑐 was used to fit 1/𝑣_depol_ vs time curves, where:

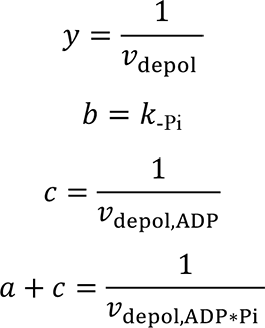

Barbed end P_i_-release rate was calculated as follows:

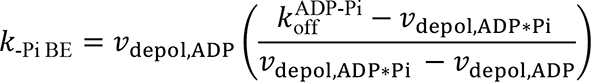

Where 𝑘_-Pi_ and 𝑘_-Pi BE_ are the phosphate release rates from the interior of the filament and the barbed end respectively. 𝑣_depol_, 𝑣_depol,ADP_ and 𝑣_depol,ADP*Pi_ are the observed depolymerization rate, depolymerization rate of ADP-actin subunits and depolymerization rate of ADP-P_i_-actin subunits respectively. ADP-P_i_-actin subunits can dissociate from the filament in two different ways; either as actin:ADP-P_i_ with a rate of 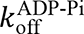 or first by releasing the Pi with a rate of 𝑘_-Pi BE_ and then dissociating with a rate of 𝑣_depol,ADP_ (ref.^20^). For our calculations we fixed 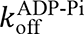 to 0.2 s^-1^ as measured previously^25^.

We estimated lower bounds for the P_i_ release rate of the N111S actin mutant by assuming the 𝑣_ADP-Pi-depol_ to either be equal or twice that of wt actin. The latter assumption was motivated by the observation that the 𝑣_depol,ADP_ rate of N111S mutant was about 2.2 times that of wt actin (Fig. 3d, Supplementary Fig. 5e).

### Sample preparation for cryo-EM

To create the short β/γ-actin filaments, 80 μM β/γ-actin:DNaseI complex was mixed with 2 μM free β/γ-actin and 25 μM mDia1_FH2_ (final concentrations) in a total volume of 50 μl G-buffer. The sample was then incubated with 5.6 μl 10x ME on ice for 1 min to exchange the ATP-associated divalent cation in actin from Ca^2+^ to Mg^2+^. Actin polymerization was induced by the addition of 6.2 μl 10x KMEH (100 mM HEPES pH 7.0, 1000 mM KCl, 20 mM MgCl_2_, 10 mM EGTA), resulting in a salt concentration of 100 mM KCl and 2 mM MgCl_2._ After incubation in a room temperature water bath for 30 seconds, we added 1 μl phalloidin (in DMSO) to the mixture to a final concentration of 80 μM. The sample was placed on ice for another 30 minutes, and was then injected onto a Superdex200increase 5/150 column (Cytiva) pre-equilibrated in 10 mM imidazole pH 7.1, 100 mM KCl, 2 mM MgCl_2_ and 1 mM EGTA on an ÄKTA Micro system (Cytiva). The eluted filament fractions were collected and directly used for cryo-EM grid preparation within 30 minutes.

The recombinant human β-actin filaments with the N111S or R183W mutation were prepared similarly as described previously for native rabbit skeletal α-actin filaments^8^. Frozen G-actin aliquots (53 μM of N111S-actin, 58 μM of R183W-actin) in G-buffer were thawed and centrifuged at 100,000*g* for 20 – 30 minutes to remove aggregates. 50 μl of G-actin was then mixed with 5.6 μl 10x ME and incubated for 5 min on ice to exchange the ATP-associated divalent cation from Ca^2+^ to Mg^2+^. Actin polymerization was induced by moving the sample to room temperature, followed by the addition of 6.2 μl 10x KMEH, resulting in a final salt concentration of 100 mM KCl and 2 mM MgCl_2._ The two mutants were polymerized 1 hour at room temperature, followed by 1 – 2 hours on ice. The long incubation period following polymerization ensured that both β-actin variants adopted the ‘aged’ ADP nucleotide state. The formed filaments were subsequently centrifuged at 200,000*g* for 30 minutes, and the F-actin pellet was resuspended in 1x KMEH (10 mM HEPES pH 7.1, 100 mM KCl, 2 mM MgCl_2_, 1 mM EGTA) supplemented with 0.02% Tween20 (v/v), to a final concentration of ∼15 μM F-actin. The resuspended material was used for cryo-EM grid preparation within an hour.

### Cryo-EM grid preparation

All cryo-EM grids were prepared through a common protocol; 2.8 µl of F-actin sample was applied to a glow-discharged R2/1 Cu 300 mesh holey-carbon grid (Quantifoil). Excess solution was blotted away and the grids were plunge frozen in liquid ethane or a liquid ethane/propane mixture using a Vitrobot Mark IV (Thermo Fisher Scientific, operated at 13 °C). The short β/γ-actin filaments, which allowed for the structure determination of the barbed end, were blotted with a blotting force of 0 for 3 seconds. The long filaments of recombinant β-actin with the N111S or R183W mutation were blotted with a blotting force of −20 for 8 seconds.

### Cryo-EM grid screening and data collection

The barbed-end dataset was collected on a 200 kV Talos Arctica cryo-microscope (Thermo Fisher Scientific) with a Falcon III direct electron detector (Thermo Fisher Scientific) operated in linear mode. Using EPU (Thermo Fisher Scientific), 1,316 movies were collected at a pixel size of 1.21 Å in 40 frames, with a total electron exposure of ∼56 e^-^/ Å^2^. The defocus values in EPU were set from −2.7 to −1.2 μm.

The datasets of N111S (9,516 movies) and R183W (7,916 movies) β-actin were collected on a 300 kV Titan Krios G3 microscope (Thermo Fisher Scientific) equipped with a K3-direct electron detector (Gatan) and a post-column BioQuantum energy filter (Gatan, slit width 15 eV). For both datasets, movies were obtained in super-resolution mode in EPU at a pixel size of 0.3475 Å in 60 frames (total electron exposure of ∼70 e^-^/Å^2^). The defocus values in EPU were set from −2.0 to −0.7 μm.

### Cryo-EM image processing

All datasets were pre-processed on-the-fly using TranSPHIRE^55^. Within TranSPHIRE, gain and beam-induced motion correction was performed using UCSF MotionCor2 v1.3.0^56^. The super-resolution mode collected data (N111S and R183W β-actin datasets) were binned twice during motion correction (resulting pixel size 0.695 Å). CTFFIND4.13^57^ was employed to estimate the contrast transfer function, and particles or filament segments were picked using SPHIRE-crYOLO^58^.

For the β/γ-actin barbed end dataset, particles were picked using SPHIRE-crYOLO in regular single particle mode^58^, resulting in 252,982 particles. The picked particles were binned 4x and extracted in a 96x96 box (pixel size 4.84 Å) in RELION 3.1.0^59^. The resulting particles.star file was then imported into CryoSPARC v.3.3.2^60^, which was used for the majority of image processing of this dataset. Importantly, the particles were processed as regular single particles without any applied helical symmetry or restraints. We first performed a 2D classification into 100 classes, thereby removing 11,275 junk particles. The remaining 241,709 particles were subjected to two rounds of heterogenous refinements. In the first round, we supplied a reference of the pointed end and a reference of a complete filament core. Through this approach we removed all particles that represented the pointed end (106,675 particles) and kept all particles that classified into the full filament. For the second round, the remaining particles (135,034 particles) were again heterogeneously refined against two references: a full filament map and a barbed end map. This allowed for the removal of 72,877 filament core particles and the isolation of 62,157 barbed end particles. These barbed end particles were then un-binned (pixel size 1.21 Å) and subjected to a non-uniform refinement, resulting in a map at a resolution of 4.07 Å. We then converted the particles to RELION (using csparc2star.py) for Bayesian polishing^61^. Afterwards, we re-imported them into CryoSPARC and ran iterative 2D classifications and 3D classifications without image alignment to remove any remaining junk particles. Following one extra round of Bayesian polishing, the final 43,618 barbed-end particles were non-uniformly refined, with the per-particle defocus estimation option switched on, to a reconstruction of 3.51-Å resolution. To further improve the density of subunits at the barbed end, we created a soft mask around the first four actin subunits (from the end) and ran a local refinement in CryoSPARC. The resulting cryo-EM density map was refined to a slightly lower resolution (3.59 Å) but showed an improved local resolution for the penultimate and ultimate subunits at the barbed end.

For the N111S and R183W β-actin datasets, filament segments were picked using the filament mode of SPHIRE-crYOLO^62^, with a box distance of 40 pixels between segments (corresponding to 27.8 Å) and a minimum number of six boxes per filament. This yielded 2,001,281 and 1,569,882 filament segments (from now on referred to as particles) for the N111S and R183W datasets, respectively, which were extracted (384x384 box) and further processed in helical SPHIREv1.4^63^. The used processing strategy was essentially the same as reported in our previous work^8^. Briefly, the extracted particles were first 2D classified using ISAC2 (refs^64, 65^) in helical mode and all non-protein picks were discarded. The particles were then refined using meridien alpha, which imposes helical restraints to limit particle shifts to the helical rise (set to 27.5 Å) to prevent particle duplication, but does not apply helical symmetry^55^. For both datasets, the first refinement was performed without mask using EMD- 15109 as initial reference, low-pass filtered to 25 Å. From the obtained reconstruction, a soft mask was created that stretched 85% of the filament length within the box (326 pixels in the Z-direction). Masked meridien alpha refinements then yielded density maps at resolutions of 2.9 Å (N111S dataset) and 3.0 Å (R183W dataset). Subsequently, the particles of both datasets were converted to be readable by RELION through sp_sphire2relion.py. In RELION, the particles were subjected to Bayesian polishing (2x) and CTF refinement, followed by a 3D classification without image alignment into 8 classes. We selected the particles that classified into high-resolution classes and removed duplicates, which yielded a final set of 1,756,928 and 1,386,604 particles for the N111S and R183W datasets, respectively. Finally, these particles were refined from local searches (sampling 0.9°) with solvent flattening Fourier shell correlations (FSCs) in RELION, yielding reconstructions at 2.30 Å (N111S) and 2.28 Å (R183W) resolution. RELION was also used to estimate the local resolutions of each density map.

### Model building, refinement and analysis

The barbed-end β/γ-actin structure was modeled as β-actin. Of the 4 amino-acid substitutions between β and γ-actin, three represent N-terminal amino-acids (D2, D3 and D4 in β-actin, E2, E3 and E4 in γ-actin) that are not visible in the density map due to flexibility. Accordingly, the only other residue (V10 in β-actin, I10 in γ-actin) that is different between both isoforms was modeled as valine. To construct an initial model for human β-actin, chain C of the 2.24-Å cryo-EM structure of rabbit skeletal α-actin in the Mg^2+^-ADP state (pdb 8A2T – 94% sequence identity to human β-actin, all water molecules removed) was rigid-body fitted in the third actin- subunit from the barbed end (A_2_) in the cryo-EM density map using UCSF ChimeraX^66^. Rabbit skeletal α-actin residues were substituted with the corresponding residues of human β-actin in Coot^67^. The resulting model was then also fitted in the densities for the A_0_, A_1_ and A_3_ actin subunits. The model for the cyclic toxin phalloidin was taken from pdb 6T1Y^35^ and rigid-body fitted into the density map. The barbed structure was iteratively refined by manual model building in Coot and Phenix real-space refine^68^ with applied Ramachandran and rotamer restraints. As described in the manuscript, the model of the ultimate A_0_ actin-subunit had to be substantially altered from the starting models.

The model of recombinant β-actin with the N111S mutation was obtained through a similar approach; chain C of pdb 8A2T (including all water molecules) was fitted into the central actin subunit of the density map. After substitution of all α-actin specific amino-acids to the corresponding β-actin residues, introducing the N111S mutation, and further manual model building in Coot, the resulting model was fitted in four more actin subunits (chains A, B, D, E) in the density map. The filament was modeled as a pentamer to capture the full interaction interface of the central subunit with its four neighboring subunits. All water molecules were first manually built, inspected and adjusted in the central subunit, and were then copied to the other chains with non-crystallographic symmetry (NCS). Because the local resolution was worse at the periphery of the reconstruction, we removed water molecules that displayed poor corresponding cryo-EM density in the non-central actin chains. The model was refined in Phenix real-space refine with NCS restraints but without Ramachandran and rotamer restraints. The model of R183W-β-actin was constructed and refined in the same way, except that the N111S-β-actin structure was used starting model for model building. A summary of the refinement quality of the structures is provided in Supplementary Table 1. Figures depicting cryo-EM density maps and protein structures were prepared using ChimeraX-1.5 (ref.^66^).

### Preparation of structural models for MD simulations

All-atom models of the core and barbed end of F-actin were prepared from respectively pdb 8A2S (F-actin core in the Mg^2+^-ADP-P_i_ state, ref.^8^) and pdb 8OI6 (the F-actin barbed end structure presented in this study). Missing residues 46-48 in the D-loop were built *de novo* with MODELLER^69^. To ensure that simulations could be compared between structural states, the barbed end structure was back-mutated from human β-actin to human/rabbit α-actin using MODELLER. The missing C-terminal residues (363-375) of the ultimate subunit in the barbed end structure were modelled using the corresponding resolved residues in the adjacent subunit as template. Phalloidin molecules were removed in the barbed end model. Hydrogen atoms were added with CHARMM c43b2 (ref.^70^). The inorganic phosphate ion was modelled in the H_2_PO_4_ ^-^ state. This is consistent with a recent quantum mechanical analysis of ATP hydrolysis by F-actin^7^, and also promotes sampling of P_i_ egress events by reducing the strength of electrostatic interactions^32^. Protonation states of histidine residues were determined with ProPka^71^. Histidine 73 was modelled as a protonated methyl-histidine (meH73^+^). For the F-actin core, a separate model with a neutral methyl-histidine 73 (meH73) was also prepared. Non-histidine titratable residues were modelled in their standard state. For the actin core models, structural water molecules were kept. As the barbed end structure was determined in the ADP-bound state, we added the inorganic phosphate ion in the same position relative to ADP as seen in the actin core structure. Structural models were embedded in an orthorhombic simulation box (20 nm 10.4 nm 8.8 nm for actin core, 18.8 nm 11.6 nm 8.8 nm for barbed end), solvated with TIP3P water molecules and supplemented with Na^+^ and Cl^-^ ions to ensure electroneutrality and reach a total salt concentration of 150 mM. CHARMM, VMD^72^ and scripts obtained from CHARMM-GUI^73^ were used for model preparation. Each model was then energy-minimized for 5000 steps of steepest descent with harmonic position restraints (force constant 4184 kJ/mol/nm²) applied on backbone atoms.

### MD simulation parameters

All simulations were performed with GROMACS 2021.5 (refs.^74, 75^) patched with the February 2022 version of the *colvars* module^76^. Energetics was described with the CHARMM36m force- field^77^. Van-der-Waals forces were smoothly switched to zero between 1.0 nm and 1.2 nm. Long-range electrostatics was treated by the Particle Mesh Ewald (PME) method. The cut-off between short-range and long-range electrostatics was initially set to 1.2 nm then was adjusted automatically by GROMACS to improve simulation performance. The length of covalent bonds involving hydrogen atoms was constrained with LINCS^78^. The leapfrog integrator with a 2 fs timestep was used for molecular dynamics. An average temperature of 310 K was maintained with the v-rescale thermostat^79^ (Berendsen for equilibration^80^); an average pressure of 1 bar was maintained with the Parrinello-Rahman barostat^81^ (Berendsen for NPT equilibration). Minimized systems were equilibrated under active backbone restraints by 1 ns of NVT simulation followed by 1 ns of NPT simulation. For production simulations, backbone restraints were turned off and the filament was kept parallel to the box using a harmonic restraint on the *orientation* quaternion (force constant 10^5^ kJ/mol) as implemented in *colvars*. MD trajectories were visualized and analyzed with VMD, Pymol (Schrödinger) and MDAnalysis^82^.

### Metadynamics protocol to sample P_i_ release

To efficiently sample P_i_ release events from actin, we developed an enhanced sampling protocol based on simulating ’swarms’ of short metadynamics trajectories^32, 83^. Metadynamics applies a history-dependent potential on a user-defined collective variable to make previously visited configurations more unstable, thus enhancing the sampling. Here, we apply metadynamics on P_i_ to drive it out of the active site. For this purpose, we concurrently applied separate conventional metadynamics biases on the three Cartesian coordinates x_P_, y_P_, z_P_ of the phosphorous atom of the inorganic phosphate ion, expressed in the internal reference frame of the actin filament. The bias was applied only on the P_i_ ion of the relevant actin subunit, namely the ultimate subunit in the barbed end, and the central subunit in the F-actin core. The bias deposition frequency was set to 0.125 /ps, the bias strength to W=1 kJ/mol and the Gaussian potential standard deviation to 0.05 nm. These parameters ensure the capture of P_i_ escape events. We stress that they are not suited for the evaluation of the converged free energy profile along a P_i_ egress path, which was not the purpose of our simulations. Each swarm was composed of 50 independent 10 ns-long metadynamics simulations launched from the same structure. Swarms were analysed by visual inspection and machine learning.

### Clustering and analysis of P_i_ egress paths

To get a broad picture of the main P_i_ egress paths sampled by metadynamics, we used hierarchical clustering in path space. For this purpose, P_i_ was considered to have escaped when the distance of the P atom from its initial position in the frame of the actin filament exceeded a cut-off value of 1.4 nm, and frames posterior to escape were discarded for the rest of the analysis. We clustered escape paths as follows. First, each P_i_ egress trajectory (represented by the time-series of the x_P_, y_P_, z_P_ coordinates defined above) was smoothed using B-spline interpolation. Then, 100 equally-spaced points were extracted along the fitted splines for each trajectory, resulting in 50 discretized paths in Cartesian space per swarm. Finally, the Euclidean distance matrix between the 50 discretized paths was computed. From this matrix, we performed hierarchical clustering of P_i_ egress paths. To visualize the paths corresponding to each cluster, we first computed each cluster’s centroid path as the cluster average in discretized path space. Then, we used the closest cluster member to the centroid as this cluster’s representative path. We found that a user-defined value of 4 clusters was sufficient to uncover the main sterically accessible P_i_ egress paths from actin. Spline interpolation was performed with SciPy^84^ and clustering with scikit-learn^85^. To complement the agnostic clustering analysis, we also evaluated the fraction of paths within a swarm for which P_i_ egress took place through the R177-N111 backdoor. For this purpose, we visually inspected the individual P_i_ egress trajectories and labelled the trajectories in which escape through the R177-N111 backdoor was observed. Escape through the R177-N111 backdoor was defined as P_i_ passing in between the side chains of R177 and N111 or in their close vicinity.

### Unbiased molecular dynamics of actin core

To obtain insights into the conformational dynamics of actin, a 1.1-microsecond unbiased MD simulation of the F-actin core (meH73^+^) was performed. For the central actin subunit, a soft positional restraint (force constant 625 kJ/mol/nm^2^) was applied on the coordinates x_P_, y_P_, z_P_ of the P_i_ ion of the central subunit to keep it close to the post-hydrolysis position. We analyzed the opening state of the R177-N111 backdoor by measuring the distance between R177CZ and N111CG atoms (N111-R177 distance) along the simulation trajectory.

### Classifier model to detect Pi egress through the R177-N111 backdoor

We trained a logistic regression binary classifier to automatically detect trajectories in which P_i_ egress happens through the R177-N111 backdoor. Our classifier takes as input a featurized P_i_ egress trajectory from a metadynamics simulation, and returns 1 if this trajectory is identified as escaping through the R177-N111 backdoor, 0 otherwise. To make the model insensitive to differences in absolute Cartesian coordinates between simulations of different structures, we developed an input trajectory featurization based on the distances of P_i_ to key residues during egress. First, to deal with escape trajectories of unequal time lengths, we computed for each trajectory the longitudinal path collective variable *s* [pathCV^86^] ranging from 0 (P_i_ in the post-hydrolysis position) to 1 (P_i_ escapes). Thus, *s* provides a common rescaled time enabling the comparison of trajectories of different lengths. The pathCV calculation was performed with an in-house Python script. The λ parameter was set to 325 nm^-2^ and the reference path was obtained by B-spline interpolation of the x_P_, y_P_, z_P_ time-series, then discretized into 30 equally-spaced points. For each simulation frame of a given P_i_ egress trajectory, we then computed *s*, along with distances between the P atom of P_i_ and the CA atom of 48 residues involved in typical egress pathways detected in the clustering and/or visual analyses (residues 12-18, 71- 74, 106-116, 120, 134-138, 141, 154-161, 177, 183, 300-301, 334-338, 370, 374). Distances were turned into Boolean contact values using a distance cut-off of 0.75 nm. Then, contact values along egress paths were grouped into 4 bins according to the pathCV *s* value, and the average P_i_ contact occupancy per bin and per residue was computed. This procedure yielded a P_i_ contact map along discretized rescaled time, which has the same 48x4 size for every P_i_ egress trajectory regardless of the actual time to P_i_ escape. Each contact map was finally flattened into a column vector whose entries were used as input features for the classifier. Training of the classifier was performed using a manually labelled data set of 300 P_i_ egress trajectories, including 70 positives. This data set was randomly split into a training and test sets in 75%/25% proportion using class balancing to preserve the proportion of true positives. The classifier was trained on the training set, and the confusion matrix was evaluated on the test set. scikit-learn was used for these tasks.

### Steered MD simulations

To simulate the opening of the R177-N111 backdoor from the filament interior, we used steered molecular dynamics to drive the central actin core subunit towards the barbed-end configuration and away from the actin core configuration. For this purpose, we applied a harmonic restraint (force constant 334124 kJ/mol/nm^2^ or 800 kcal/mol/Å^2^) moving with constant velocity on the ΔRMSD, *i.e.,* the difference in RMSD between actin core and barbed end structures. The biasing potential 𝑈 acting on atomic configuration 𝑥 at time 𝑡 was:

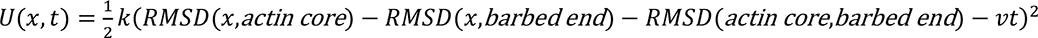

with v = 0.0022 nm/ns and where *RMSD(x, actin core)* [respectively *RMSD(x, barbed end)*] refers to the RMSD of atomic configuration *x* with respect to the actin core structure (respectively the barbed end structure). RMSD values were computed on the CA atoms of residues 8-21, 29-32, 35-38, 53-68, 71-76, 79-93, 103-126, 131-145, 150-166, 169-178, 182-196, 203-216, 223-232, 238-242, 246-250, 252-262, 274-284, 287-295, 297-300, 308-320, 326-332, 338-348, 350-355, 359-365, 367-373. To mimic the influence of a long actin filament in cellular conditions, static RMSD restraints (force constant 4184 kJ/mol/nm^2^) were applied separately on the CA atoms of two pairs of lateral actin subunits flanking the central subunit. To decouple backdoor opening from P_i_ movement, a positional harmonic restraint was applied on the P atom Cartesian coordinates x_P_, y_P_, z_P_ (force constant 625 kJ/mol/nm^2^). Forty separate SMD simulations were run (20 with meH73^+^ and 20 with meH73). The following structural observables were computed along each trajectory: N111-R177 distance (see above), sensor/Pro-rich loops distance (defined as the distance between the centers of geometry of CA atoms of sensor loop residues 70-75, and CA atoms of Pro-rich loop residues 105-115), H161- Q137 distance (defined as the distance between H161NE2 and Q137OE1), and H161 side chain torsion angle 𝜒_2_. To evaluate the capacity of structures sampled by SMD to release P_i_ through the R177-N111 backdoor, frames were extracted at t = 0 ns (equilibrated structure), t = 15 ns, 25 ns, 35 ns, 45 ns, 50 ns, 65 ns, 75 ns, 85 ns and 100 ns. Then, each of these frames was used to launch a swarm of 50 metadynamics simulations to probe P_i_ release according to the protocol described above. A static ΔRMSD restraint with same force constant and reference value as the SMD potential was applied to preserve the configuration of the actin core subunit. For each swarm, all trajectories were analyzed with the logistic regression classifier. Finally, the proportion p_BD_ of trajectories with predicted R177-N111 backdoor escape was computed for each swarm.

## Acknowledgements

We thank S. Bergbrede for the purification of β-actin mutants and overall wet lab support; J. Funk and F. Merino for initial experiments; and M. Boiero Sanders and R.S. Goody for valuable discussions and critical proofreading of the manuscript. This work was supported by funds from the Max Planck Society (to G.H. and S.R.) and the European Research Council under the European Union’s Horizon 2020 Programme (ERC-2019-SyG, grant no. 856118 to S.R). A.B. is supported by an EMBO long-term fellowship. W.O. is supported by a postdoctoral fellowship from the Alexander von Humboldt foundation. P.B. thanks P. Bastiaens for continuous support.

## Author contributions

W.O., G.H., P.B. and S.R. conceived the project. A.R., P.B., and W.O. performed protein purifications and biochemical assays. W.O. and P.B. prepared samples for cryo-EM. W.O. and O.H. collected and W.O. processed cryo-EM data and built the protein models. F.E.C.B. carried out all molecular dynamics simulations. A.B. performed the yeast drop test assays. G.H., P.B. and S.R. supervised the project. W.O., F.E.C.B., P.B. and S.R. wrote the manuscript, with critical input from all authors.

## Additional Information

**Correspondence and requests for materials** should be addressed to G.H, P.B. or S.R.

## Data availability

The cryo-EM maps generated in this study have been deposited in the Electron Microscopy Data Bank (EMDB) under accession codes (dataset in brackets): EMD-16887 (β/γ- actin barbed end), EMD-16888 (R183W-F-actin) and EMD-16889 (N111S-F-actin). These depositions include sharpened and unsharpened maps, unfiltered half-maps and the masks used for refinements. The associated protein models have been deposited in the Protein Data Bank (PDB) with accession codes 8OI6 (β/γ-actin barbed end), 8OI8 (R183W-F-actin) and 8OID (N111S-F-actin). The following previously published protein models were used for data analysis and comparisons: 8A2S [https://doi.org/10.2210/pdb8A2S/pdb], 8A2T [https://doi.org/10.2210/pdb8A2T/pdb] and 2V52 [https://doi.org/10.2210/pdb2V52/pdb]. We used EMD-15109 [https://www.ebi.ac.uk/emdb/EMD-15109] as 3D model for the first refinements of R183W- and N111S-F-actin. All scripts used for data analysis of single-filament assays can be retrieved from https://github.com/iamankitroy/Actin-Pi-Release. MD simulation models and protocols, MD simulation datasets and Jupyter notebooks to reproduce the analyses reported in Fig. 2 and Supplementary Figs. 4, 10, 12 have been deposited in Zenodo (10.5281/zenodo.7765025). All other materials are available from the corresponding authors upon request.

