## Supplementary Figures for "Molecular mechanisms of inorganic-phosphate release from the core and barbed end of actin filaments"

**Supplementary Table 1. Cryo-EM data collection, refinement and validation statistics.**

| dataset | **β/γ-actin barbed end**<br>EMD-16887<br>PDB 8OI6 | **β-actin-R183W**<br>EMD-16888<br>PDB 8OI8 | **β-actin-N111S**<br>EMD-16889<br>PDB 8OID |
|---|---|---|---|
| **Data collection and processing** | | | |
| Magnification | 120,000 | 130,000 | 130,000 |
| Voltage (kV) | 200 | 300 | 300 |
| Electron exposure (e–/Å$^2$) | 56 | 70 | 69 |
| Defocus range (μm) | -1.2 to -2.7 | -0.7 to -2.0 | -0.7 to -2.0 |
| Pixel size (Å) | 1.21 | 0.695 | 0.695 |
| Symmetry imposed | C1 | C1 | C1 |
| Initial particle images (no.) | 252,982 | 1,569,882 | 2,001,281 |
| Final particle images (no.) | 43,618 | 1,286,604 | 1,756,928 |
| Map resolution (Å)<br>0.143 FSC threshold | 3.59 | 2.28 | 2.30 |
| Map resolution range (Å) | 3.5 – 5.5 | 2.2 – 3.7 | 2.2 – 3.9 |
| Measured helical symmetry[1] | | | |
| Helical rise (Å) | 27.47 | 27.58±0.08 | 27.58±0.05 |
| Helical twist (°) | -165.7 | -166.5±0.1 | -166.5±0.3 |
| **Refinement** | | | |
| Initial model used (PDB code) | 8A2T | 8OID | 8A2T |
| Model resolution (Å)<br>0.5 FSC threshold | 3.80 | 2.30 | 2.30 |
| Map sharpening *B* factor (Å$^2$) | -140 | -55 | -60 |
| Model composition | | | |
| Non-hydrogen atoms | 11,573 | 14,895 | 15,206 |
| Protein residues | 1,449 | 1,815 | 1,850 |
| Waters | 0 | 585 | 616 |
| Ligands | 8 | 10 | 10 |
| *B* factors (Å$^2$) | | | |
| Protein | 80.55 | 32.06 | 31.10 |
| Waters | - | 27.36 | 27.47 |
| Ligand | 73.66 | 24.35 | 21.49 |
| R.m.s. deviations | | | |
| Bond lengths (Å) | 0.002 | 0.004 | 0.004 |
| Bond angles (°) | 0.596 | 0.568 | 0.597 |
| Validation | | | |
| EM-ringer score | 1.83 | 5.56 | 5.49 |
| MolProbity score | 1.40 | 1.13 | 1.45 |
| Clashscore | 7.23 | 3.42 | 4.07 |
| Poor rotamers (%) | 0.00 | 0.65 | 0.96 |
| Ramachandran plot | | | |
| Favored (%) | 98.31 | 98.60 | 96.16 |
| Allowed (%) | 1.62 | 1.40 | 3.84 |
| Disallowed (%) | 0.07 | 0 | 0 |

[1]For the β/γ-actin barbed end structure, the helical rise and twist between the ultimate ($A_0$) and penultimate ($A_1$) subunit are reported. For the filament structures, the helical parameters were estimated from the atomic model of five consecutive subunits independently fitted to the map as described previously (ref. 8).

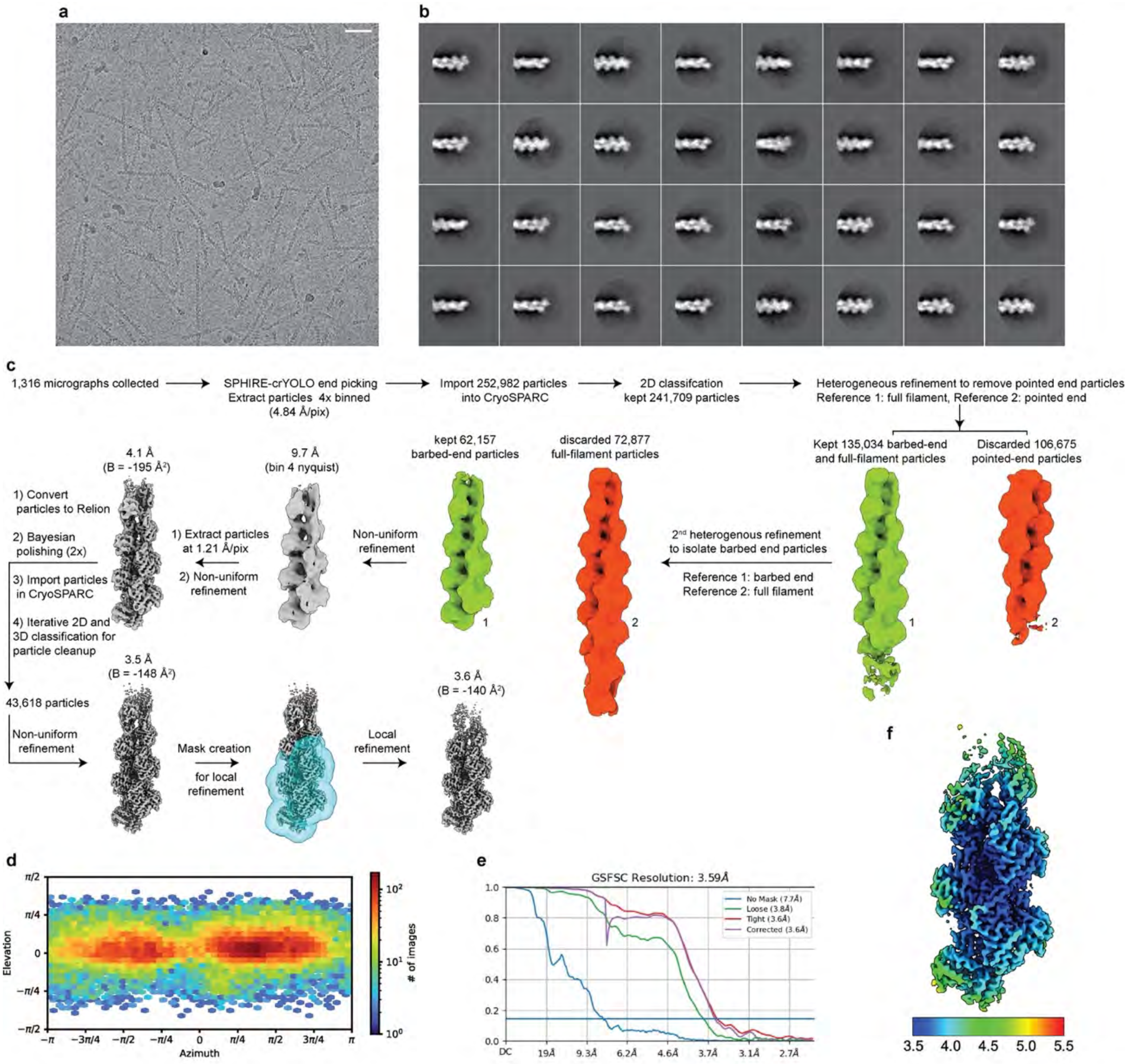


**Supplementary Fig. 1 Cryo-EM image-processing of the F-actin barbed end. a** Micrograph depicting short β/γ-actin filaments frozen in vitreous ice, at a defocus of -2.2 µm. The shown micrograph is an example image from a total dataset of 1,316 micrographs. The scale bar is 400 Å. **b** Exemplary 2D-class averages of the F-actin barbed end reconstruction, generated by RELION. The box size is 465 × 465 Å$^2$. **c** Image processing strategy that was employed to determine the β/γ-actin barbed end structure. All maps are shown in the same orientation. **d** Angular distribution of the barbed end particles used to reconstruct the final cryo-EM map, generated by CryoSPARC. **e** Fourier-shell correlation plots for the barbed structure of gold-standard refined half-maps, computed by CryoSPARC. The FSC=0.143 threshold is annotated. **f** Local-resolution estimation of the barbed end reconstruction, computed through RELION.

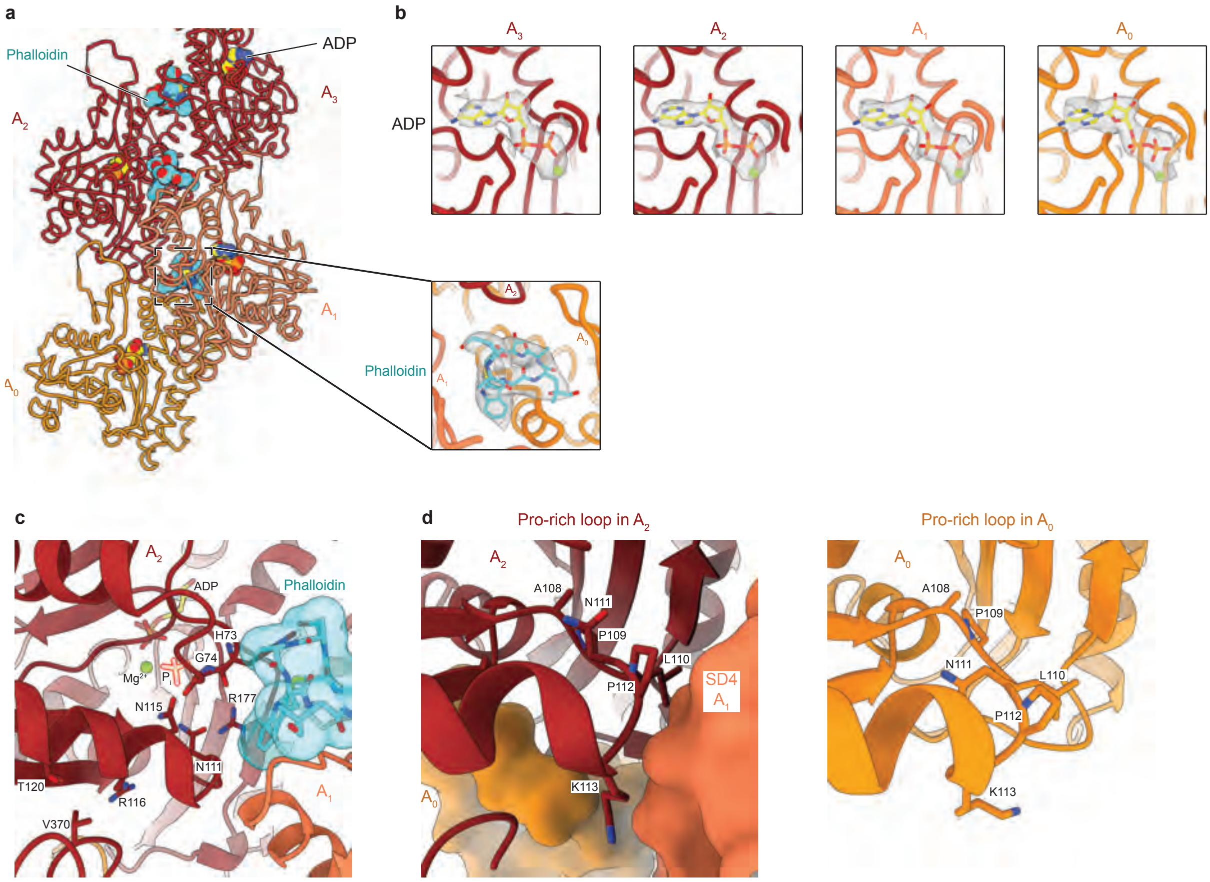


**Supplementary Fig. 2. Small-molecule binding sites and Pro-rich loop arrangement in the F-actin barbed end structure. a** Simplified cartoon representation of the barbed end structure. Actin subunits are annotated. ADP and phalloidin are shown as spheres with carbon atoms colored cyan and yellow, respectively. **b** Cryo-EM densities of the ADP and phalloidin binding sites with fitted models. F-actin is shown as simplified cartoon. **c** Zoom of the phalloidin-binding site near the R177-N111 backdoor in actin subunit $A_2$. Phalloidin is shown in stick representation with a semi-transparent surface. Residues of the proposed backdoor (R177, N111, H73, G74) are annotated, as well as other residues (N115/R116, T120/V370) that may represent other $P_i$ escape sites. Phalloidin would only interfere with disruption of the classical backdoor. **d** Arrangement of the Pro-rich loop in subunits $A_2$ (left panel) and $A_0$ (right panel) in the barbed end structure. In subunit $A_2$, the Pro-rich loop conformation is stabilized by SD4 of subunit $A_1$. Conversely, the Pro-rich loop of subunit $A_0$ does not interact with other subunits.

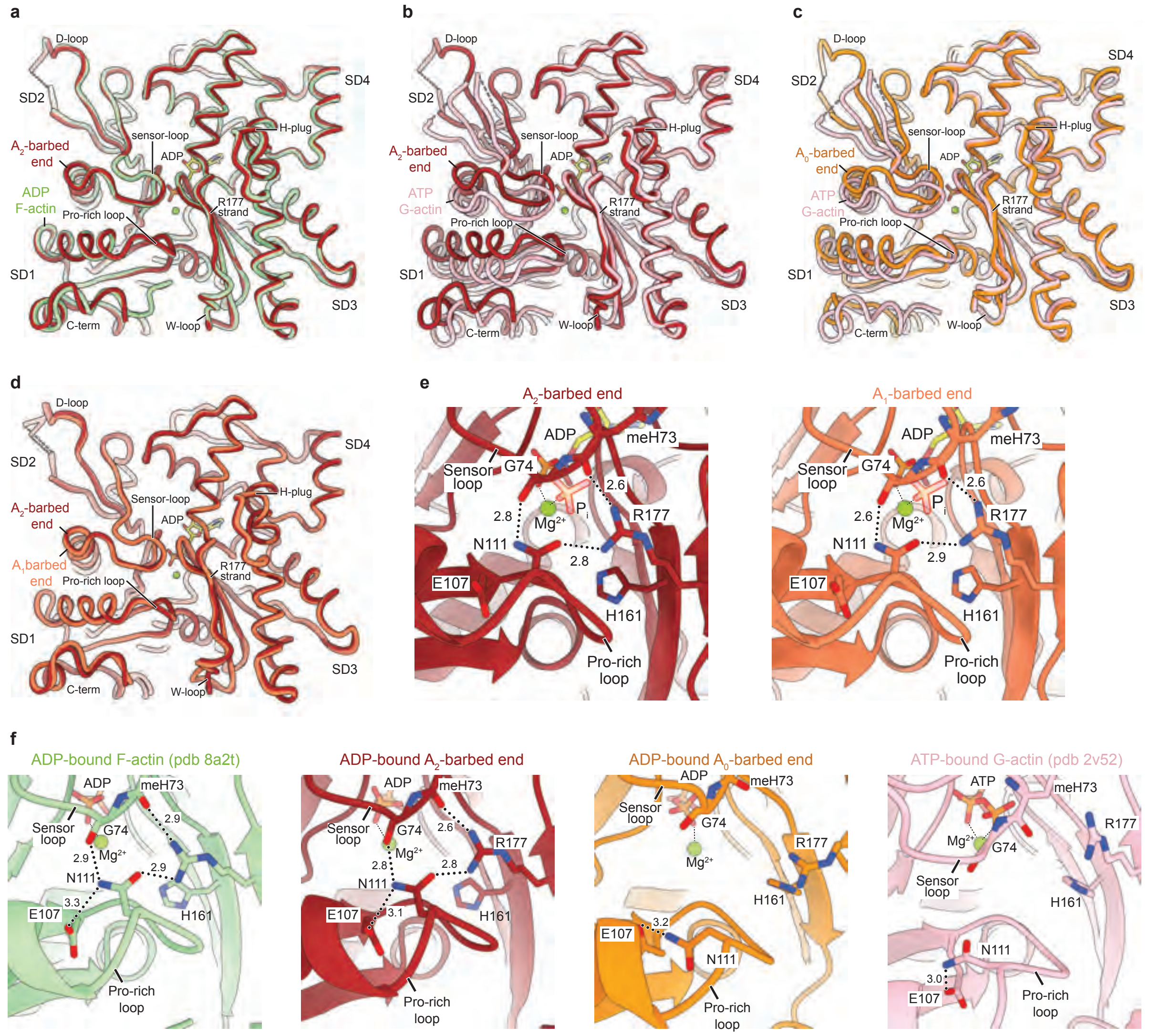


**Supplementary Fig. 3. Alignment of barbed-end subunit structures with previously determined actin structures. a** Superimposition of the $A_2$ subunit of the barbed end structure (colored dark-red) with the structure of ADP-bound filamentous α-actin (pdb 8a2t, colored green). **b** Superimposition of the $A_2$ subunit of the barbed end structure with the structure of ATP-bound monomeric α-actin (pdb 2v52, colored pink). **c** Superimposition of the $A_0$ subunit of the barbed end structure (colored orange) with the structure of ATP-bound monomeric α-actin (pdb 2v52). **d** Superimposition of the $A_2$ (colored dark-red) and $A_1$ (colored orange-red) subunits of the barbed end structure. The only observed change in penultimate subunit $A_1$ is a small rearrangement of the W-loop. This change in the W-loop is expected, because this region would interact with the D-loop of the missing subunit. **e** Zoom of the R177-N111S backdoor as seen from the F-actin exterior for the $A_2$ and $A_1$ subunits of the barbed end structure. Amino acids that form the backdoor are annotated. Phalloidin is hidden for clarity. Although the subunits adopt the ADP state, $P_i$ from pdb 8a2s (F-actin in the $Mg^{2+}$-ADP-$P_i$ state) is shown semi-transparently to emphasize the $P_i$-binding site. **f** Architecture of the $P_i$-release backdoor in structures of ADP-bound F-actin (left), the ADP-bound $A_2$ subunit at the F-actin barbed end (middle-left), the ADP-bound $A_0$ subunit at the F-actin barbed end (middle-right) and ATP-bound G-actin (right).

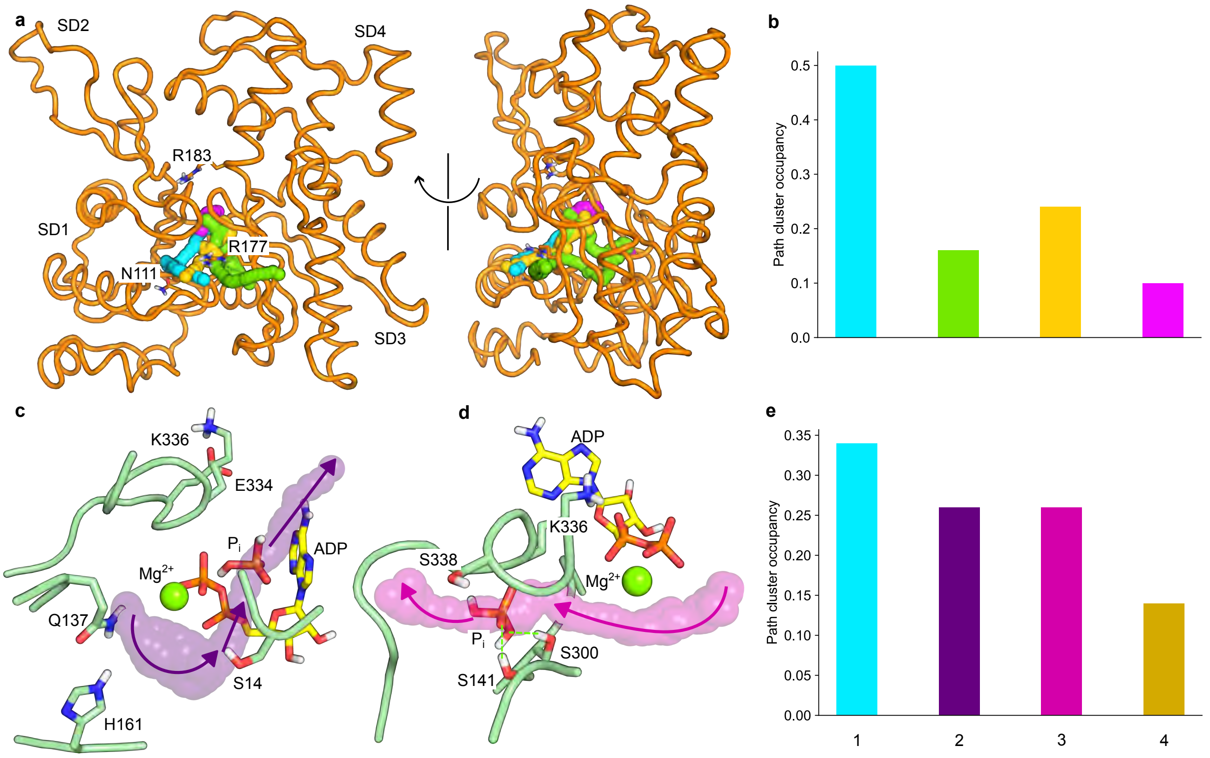


**Supplementary Fig. 4. Enhanced sampling simulations of $P_i$ release from the F-actin barbed end and core. a** Representative egress paths from the barbed end show a clear preference for egress through the open R177-N111 backdoor. **b** Path cluster occupancies in barbed end simulations with the same color code as in panel **a**. **c** A representative path of $P_i$ egress through pathway 2 (purple) in F-actin core simulations. The simulation revealed a highly bent ADP conformer that allowed $P_i$ to escape. Such a drastic rearrangement of the nucleotide binding pocket is physically implausible. In addition, this egress path would not be directly affected by phalloidin and jasplakinolide binding. Hence, this pathway was not considered for further experimental validation. **d** A representative path of $P_i$ egress through pathway 3 (magenta) in F-actin core simulations. This escape would also not be directly affected by phalloidin and jasplakinolide binding. Hence, it was not considered for further experimental validation. **e** Path cluster occupancies in actin core simulations with the same color code as in Fig. 2. Pathways 1 and 4 are further discussed in the main text.

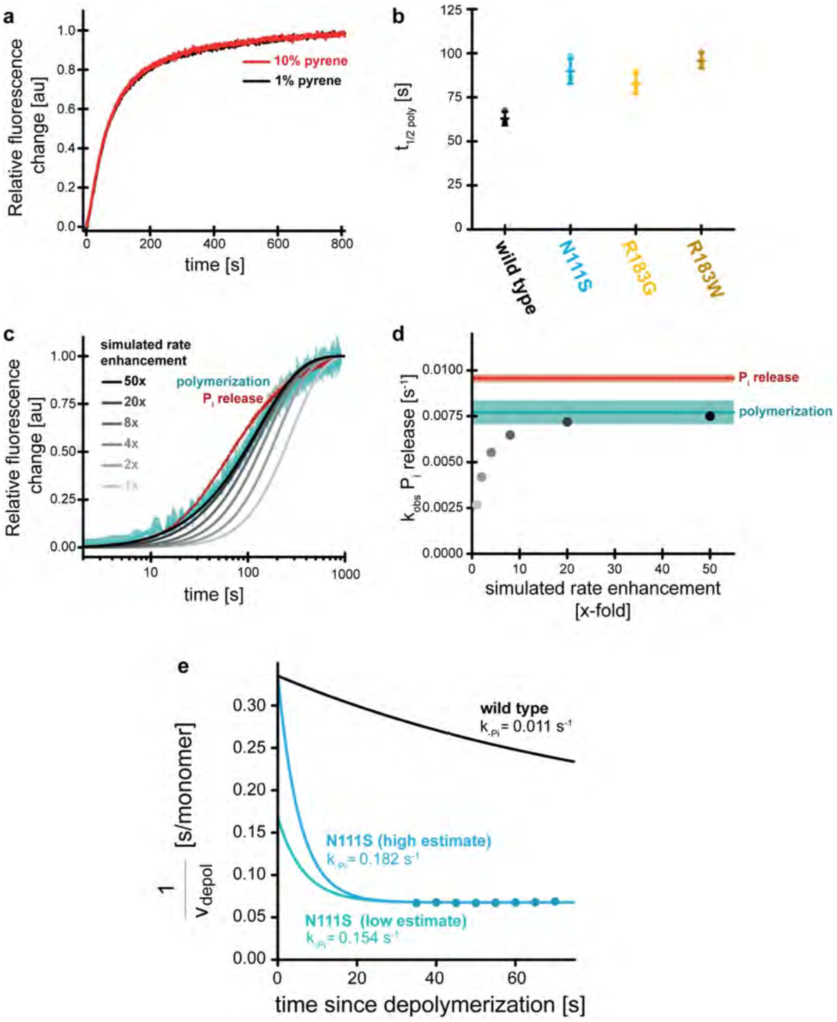


**Supplementary Fig. 5 Control experiments and kinetic analysis of $P_i$ release from actin filaments.** **a** Time courses of the normalized fluorescence intensity of 30 µM MDCC-PBP ($P_i$ sensor) from 10 µM N111S actin containing either 1% (black) or 10% (red) wild-type, pyrene α-actin, seeded with 100 nM spectrin-actin seeds after initiation of polymerization (t=0s). **b** Characteristic half-times of polymerization for indicated actin variants as determined from mono-exponential fits to the observed polymerization time-courses (Fig. 3b). **c** Semi-logarithmic plot of simulated $P_i$ release reaction kinetics (gray to black) depending on the enhancement of $P_i$ release rate constant (x-fold over wild-type actin as indicated) compared to the observed time courses of polymerization (cyan) and of $P_i$ release (red) for N111S actin. **d** Apparent rates of $P_i$ release, obtained from mono-exponential fits of the simulated data shown in c as a function of the enhancement of $P_i$ release rate constant (x-fold over wild-type actin as indicated). The cyan and red areas indicate the observed apparent rates of polymerization (cyan) and $P_i$ release (red) with dark colors being the average and the light colors representing SD. Note that an enhancement of the $P_i$ release rate constant by at least 15-fold is required for the apparent rate of $P_i$ release to fall within the error margin of the observed polymerization rate. **e** Estimation of lower bounds for the phosphate release rate of N111S mutant. Rates were calculated by fitting exponential decay functions to the observed depolymerization velocity data, so that the observed velocities converged to $v_{depol,ADP}$ and the $v_{depol,ADP-Pi}$ rate was either assumed to be equal (high estimate, blue) or twice that of the wild type (low estimate, turquoise). The latter assumption was motivated by the observation that the $v_{depol,ADP}$ rate of N111S mutant was about 2.2 times that of wild-type actin (Fig. 5d).

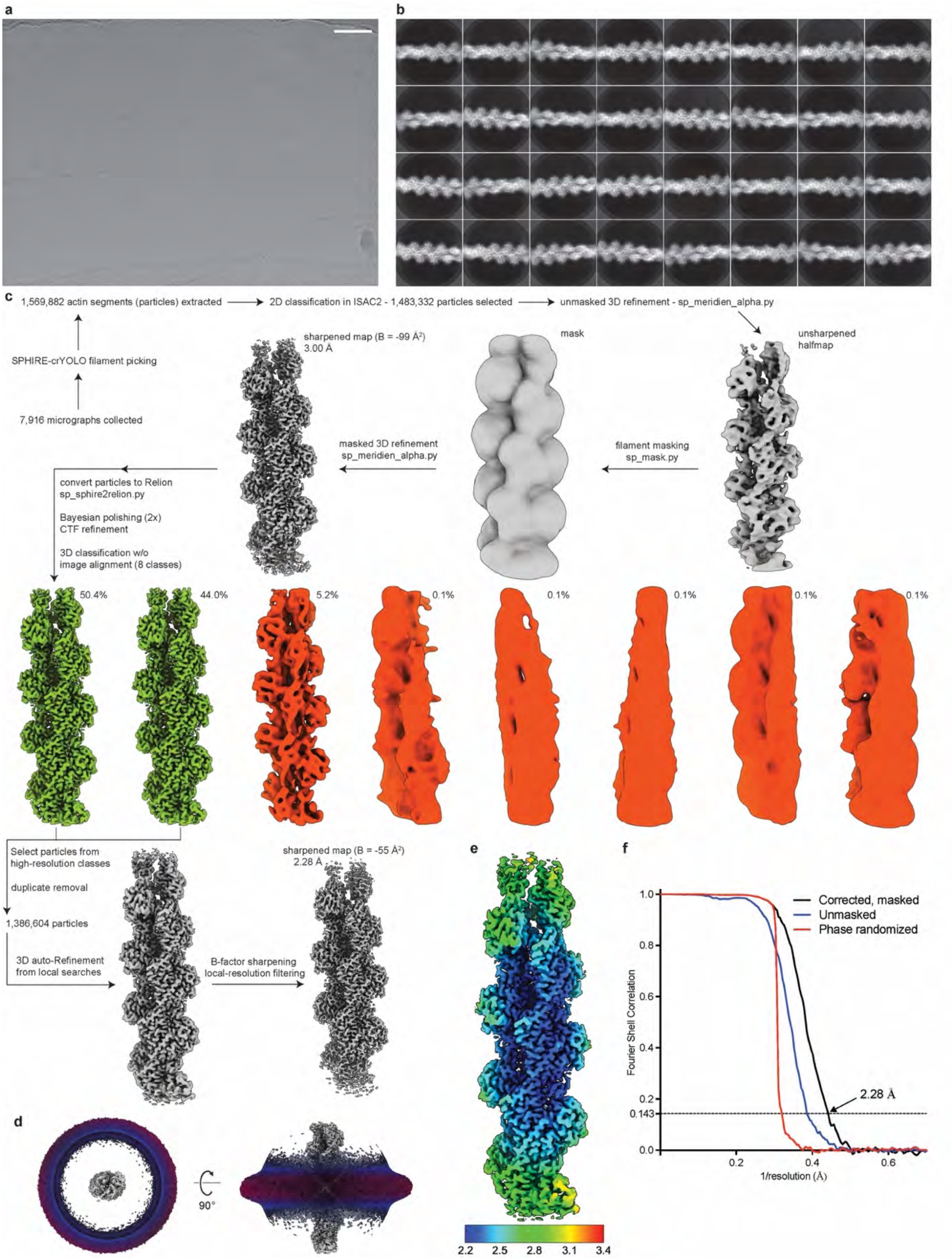


**Supplementary Fig. 6 Processing of the R183W-F-actin dataset. a** Micrograph depicting R183W-F-actin frozen in vitreous ice, at a defocus of -2.3 µm. The shown micrograph is an example image from a total dataset of 7,916 micrographs. The scale bar is 400 Å. **b** Exemplary 2D-class averages of the R183W-F-actin particles, computed through RELION. The box size is 267 × 267 Å$^2$. **c** Image processing strategy that was employed to determine the R183W-F-actin structure. All maps are shown in the same orientation. **d** Angular distribution of the barbed end particles used to reconstruct the final cryo-EM map, shown along the filament axis (left) and orthogonal to the filament axis (right). **e** Local-resolution estimation of the R183W-F-actin density map, calculated by RELION. The bar depicts local resolution in Å. **f** Fourier-shell correlation plots for gold-standard refined masked (black), unmasked (blue) and high-resolution phase randomized (red) half-maps of R183W-F-actin particles. The FSC = 0.143 threshold is shown as a dashed line.

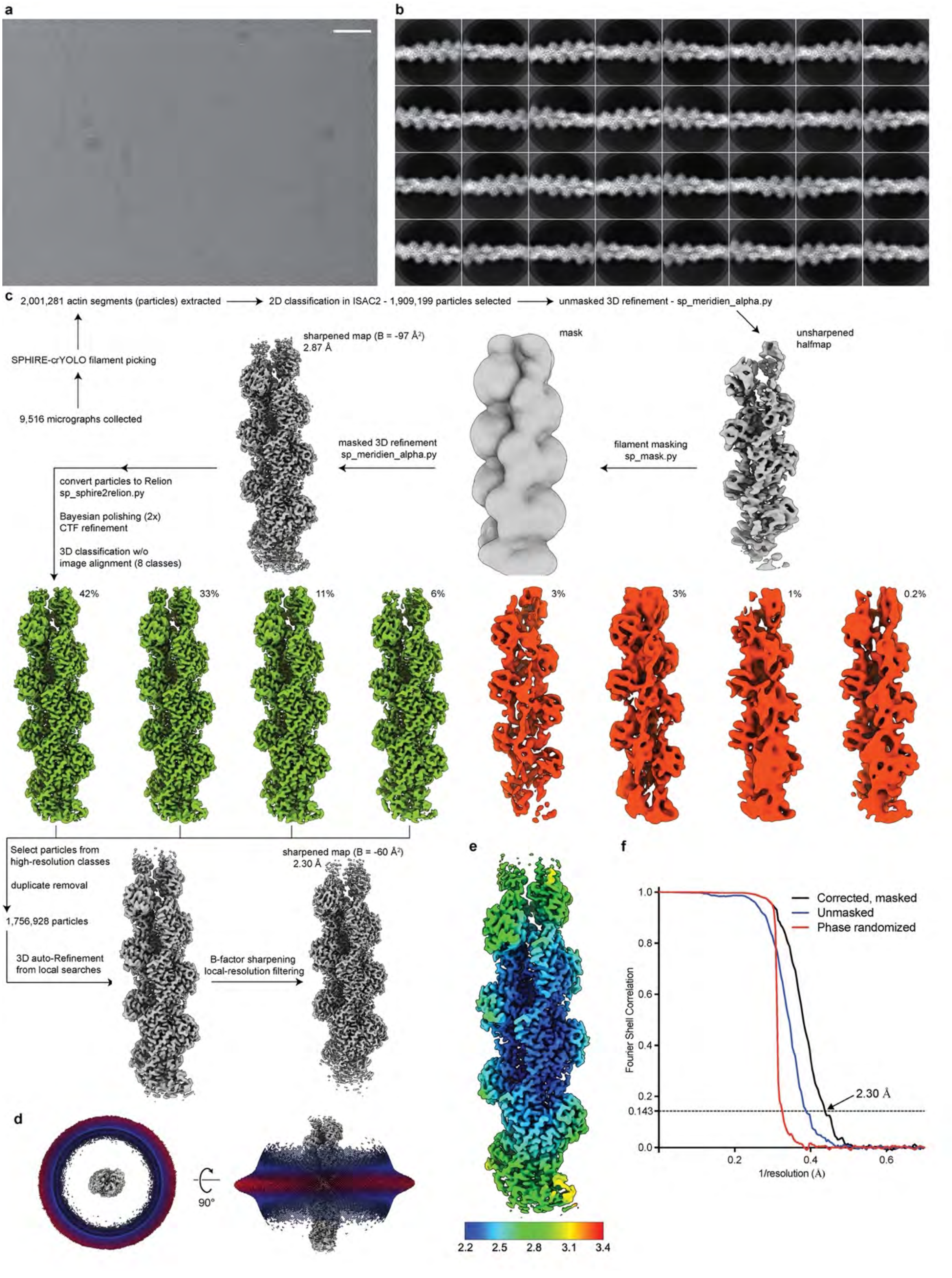


**Supplementary Fig. 7: Processing of the N111S-F-actin dataset. a** Micrograph depicting N111S-F-actin frozen in vitreous ice, at a defocus of -1.9 µm. The shown micrograph is an example image from a total dataset of 9,516 micrographs. The scale bar is 400 Å. **b** Exemplary 2D-class averages of the N111S-F-actin particles, computed through RELION. The box size is

267 × 267 Å$^2$. **c** Image processing strategy that was employed to determine the N111S-F-actin structure. All maps are shown in the same orientation. **d** Angular distribution of the barbed end particles used to reconstruct the final cryo-EM map, shown along the filament axis (left) and orthogonal to the filament axis (right). **e** Local-resolution estimation of the N111S-F-actin density map, calculated by RELION. The bar depicts local resolution in Å. **f** Fourier-shell correlation plots for gold-standard refined masked (black), unmasked (blue) and high-resolution phase randomized (red) half-maps of N111S-F-actin particles. The FSC = 0.143 threshold is shown as a dashed line.

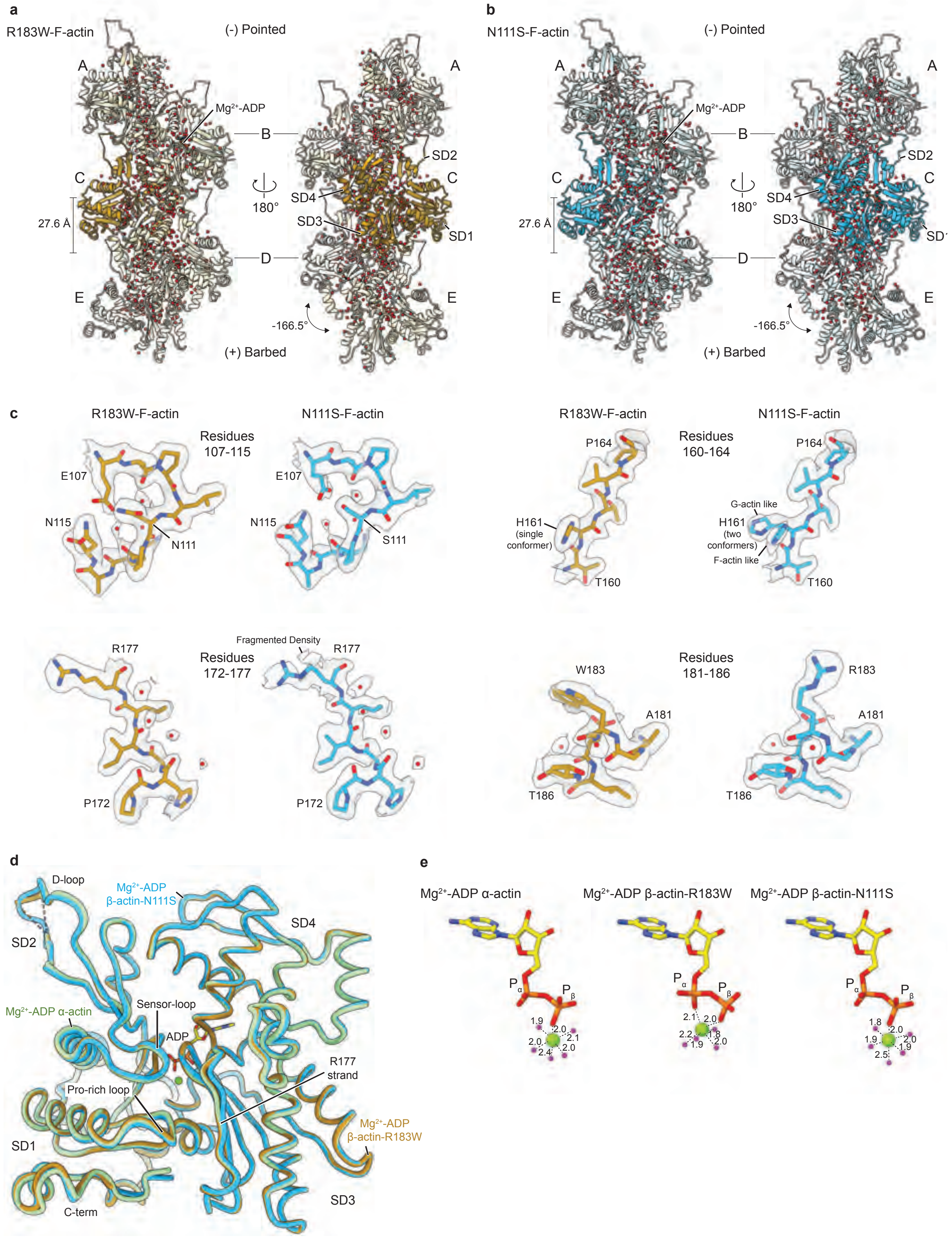


**Supplementary Fig. 8. High-resolution structures of β-actin variants R183W and N111S.** **a, b** Structures of R183W-F-actin **(a)** and N111S-F-actin **(b)** shown as cartoon orthogonal to the filament axis. The central subunits are colored gold and blue, respectively. Water molecules modeled in both structures are depicted as red spheres. The pointed and barbed end directions are annotated, as well as the helical rise and twist of both mutant structures. **c** Cryo-EM densities and fitted atomic models for selected regions of both reconstructions. Specifically, amino-acid environments near the mutated residues are shown to highlight differences between both structures. **d** Alignment of single subunits of filamentous wild-type α-actin (pdb 8a2t, colored green), R183W-β-actin (gold) and N111S-β-actin (blue) in the $Mg^{2+}$-ADP state. The F-actin subdomains and regions important for $P_i$ release are annotated. **e** Nucleotide arrangement in high-resolution $Mg^{2+}$-ADP-bound F-actin structures of wild-type α-actin (pdb 8a2t, left), R183W-β-actin (middle) and N111S-β-actin (right). The arrangements in wild-type α-actin and N111S-β-actin are similar, whereas the $Mg^{2+}$ ion adopts a different position in the R183W-β-actin structure.

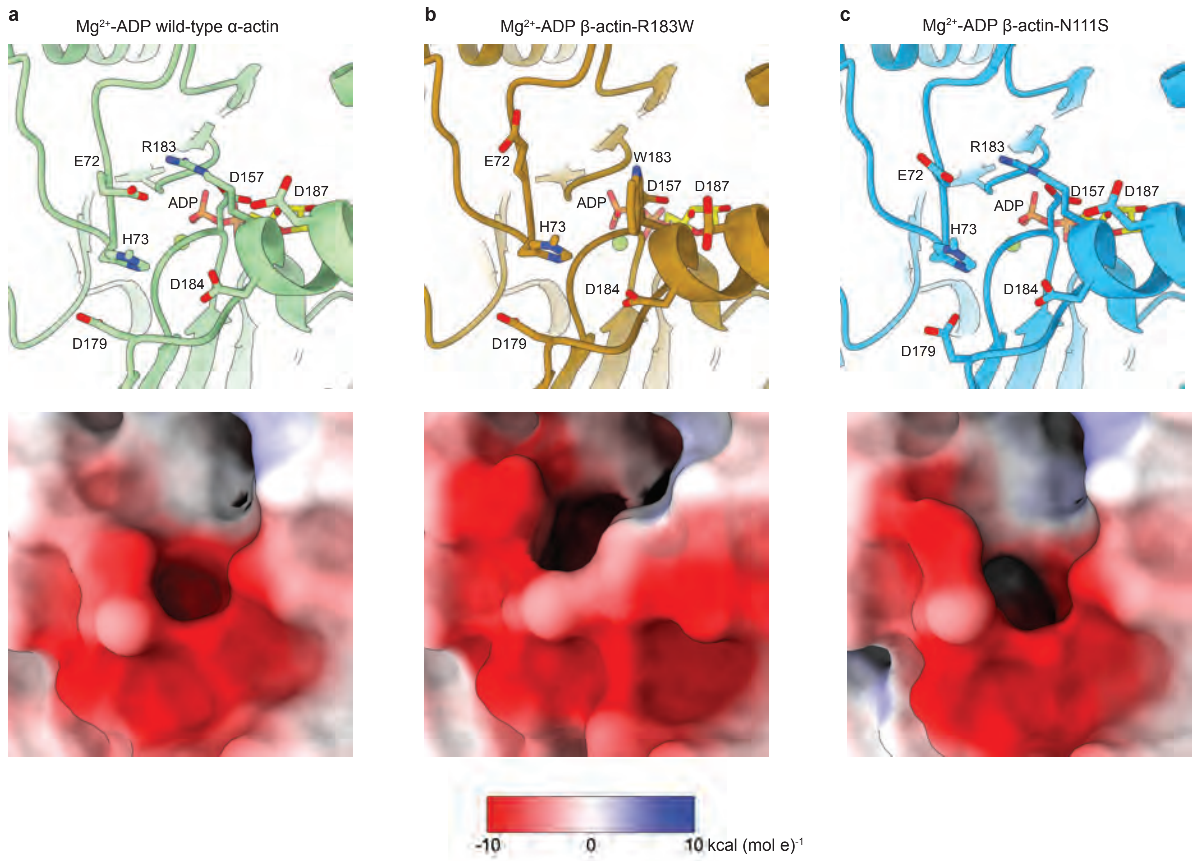


**Supplementary Fig. 9. Acidic amino-acid environment around residue 183. a-c** Arrangement near residue 183, as show from the filament exterior for filamentous wild-type α-actin **(a)** (pdb 8a2t, colored green), R183W-β-actin (gold) **(b)** and N111S-β-actin (blue) **(c)** in the $Mg^{2+}$-ADP state. The top image depicts F-actin as cartoon and charged amino acids near residue-183 as cartoon and sticks. In the lower panel, F-actin is shown as surface, which is colored by electrostatic Coulomb potential ranging from −10 kcal $(mol\ e)^{-1}$ (red) to +10 kcal $(mol\ e)^{-1}$ (blue). The surface is more negatively charged in the R183W-F-actin structure.

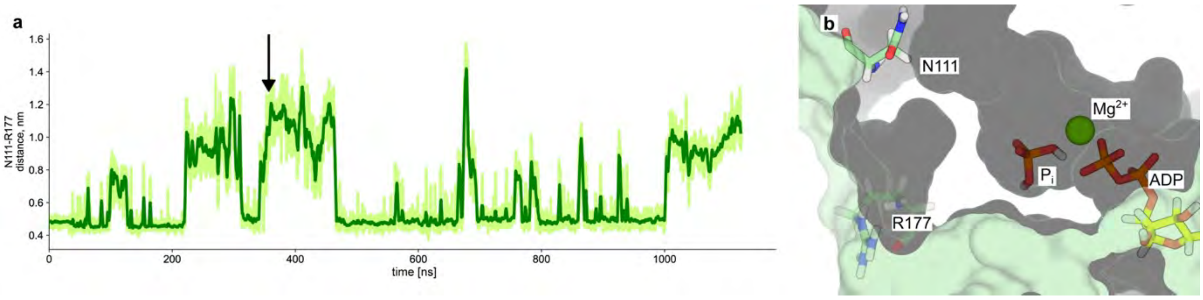


**Supplementary Fig. 10. The R177-N111 hydrogen bond reversibly breaks and re-forms in unbiased MD simulations. a** Time-series of the R177CZ-N111CG distance, depicted in light green. A distance larger than 0.6 nm indicates disruption of the hydrogen bond. For clarity, a centered 2 ns-moving average is shown in dark green. The arrow indicates the used time-frame to capture the F-actin conformation shown in panel b. **b** Example configuration with a broken R177-N111 hydrogen bond sampled at 360 ns during MD simulation. The surface representation suggests that the backdoor is closed despite the broken R177-N111 hydrogen bond. Solvent accessible area for $P_i$ is depicted in dark grey. There is no accessible path for $P_i$ to escape from the F-actin interior.

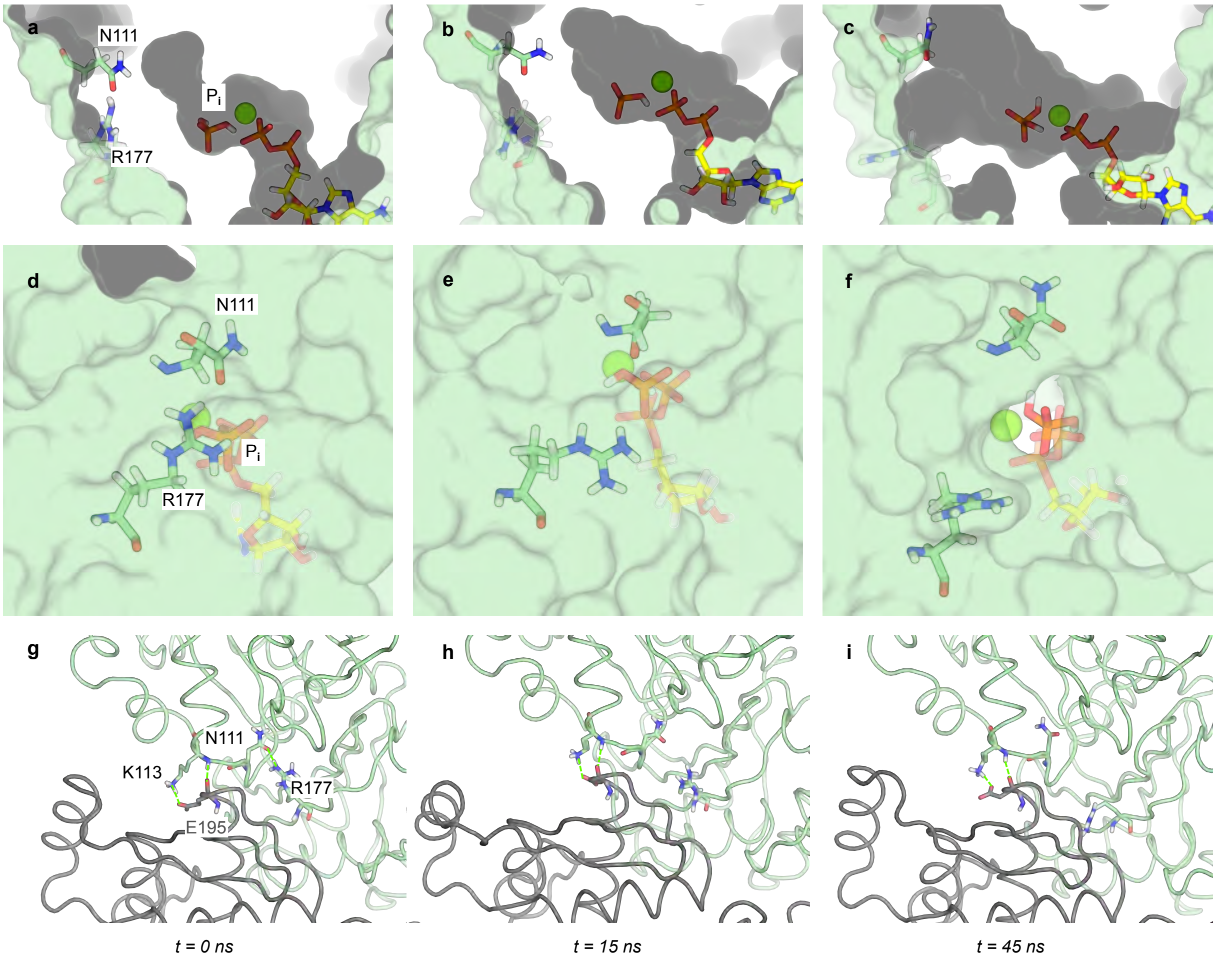


**Supplementary Fig. 11 Snapshots from SMD simulation (neutral meH73, replicate 1).** Frames are the same as in Fig. 6. **a-c** Solvent-accessible surface representation of the nucleotide binding site and R177-N111 backdoor, side view. The solvent-accessible volume is shown in dark grey. **d-f** Solvent-accessible surface representation of the R177-N111 backdoor, external view. Note that in **f** Pi is visible from the outside. **g-i** Close-up on the inter-subunit contacts.

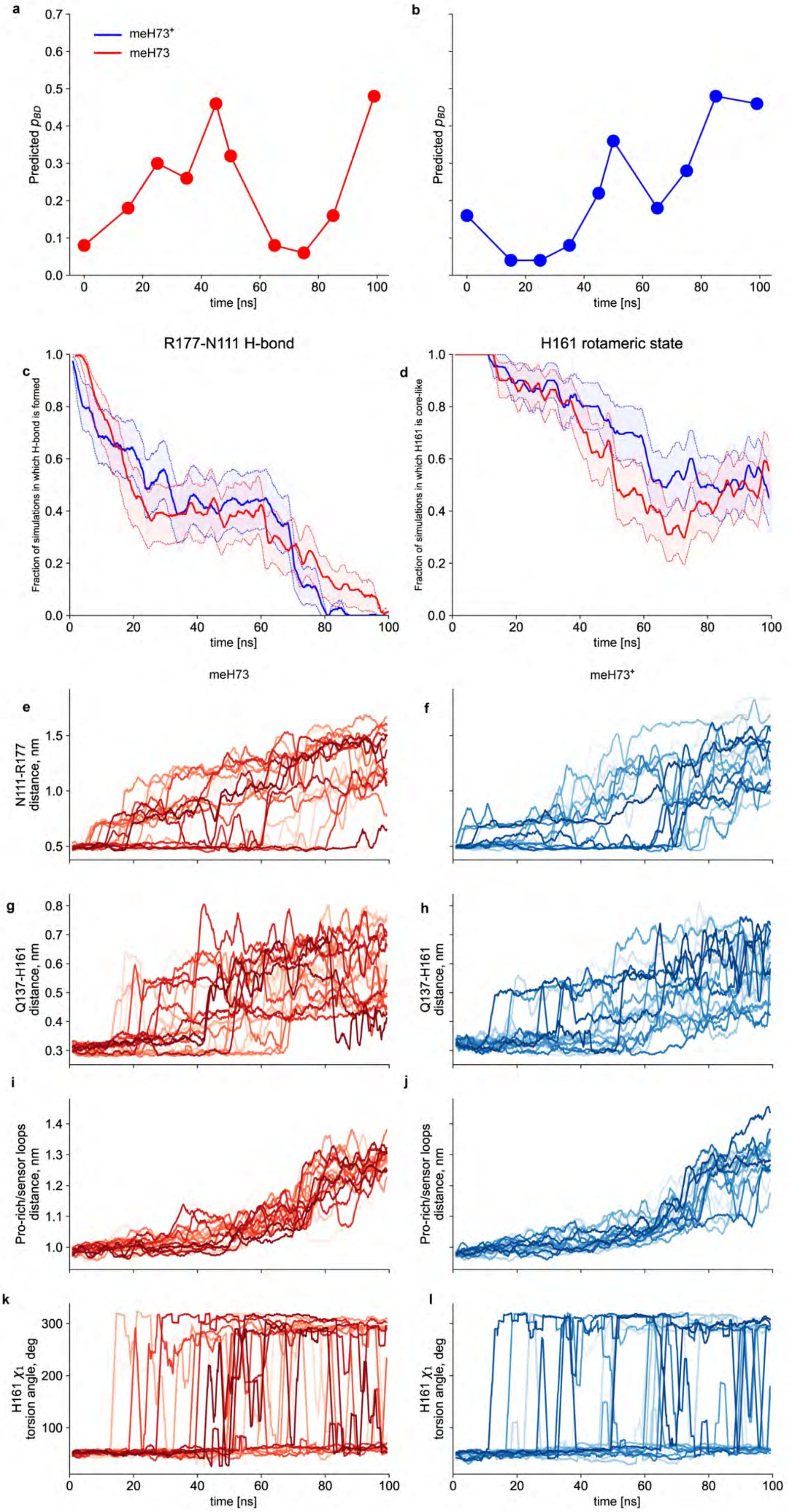

a
b
meH73⁺
meH73
Predicted $p_{BD}$
time [ns]
R177-N111 H-bond
H161 rotameric state
c
d
Fraction of simulations in which H-bond is formed
Fraction of simulations in which H161 is core-like
meH73
meH73⁺
e
f
N111-R177 distance, nm
g
h
Q137-H161 distance, nm
i
j
Pro-rich/sensor loops distance, nm
k
l
H161 $\chi_1$ torsion angle, deg

**Supplementary Fig. 12 Evolution of key observables during SMD simulations. a** Predicted fraction $_{pBD}$ of $P_i$ egress through the R177-N111 backdoor along SMD replicate 1, with neutral meH73, as obtained by classifier analysis of the enhanced sampling trajectories. **b.** Predicted fraction $_{pBD}$ of $P_i$ egress through the R177-N111 backdoor along SMD replicate 1, with positively charged meHis73, as obtained by classifier analysis of the enhanced sampling trajectories. **c**. Survival analysis of the R177-N111 hydrogen bond in SMD simulations. For each protonation state of meH73, we computed the fraction of trajectories with the R177-N111 hydrogen bond still formed at time t. A distance cut-off of 0.6 nm between atoms CZ of R177 and CG of N111 was used to define a formed hydrogen bond. The solid lines represent the 2 ns-rolling average of the survival fraction. The dotted lines represent the 2 ns-rolling averages of the survival fraction ± SEM. The background overlays represent the raw values for the survival fraction ± SEM. **d.** Survival analysis of the rotameric state of H161. For each protonation state of meH73, we computed the fraction of trajectories with H161 in a core-like rotameric state. H161 side-chain rotameric states were defined by $\chi_1$ torsion angle being above or below 180 deg. Representation conventions are the same as for panel **c**. **e-i.** Evolution of structural observables during for all SMD simulation replicates. For clarity, 2 ns-rolling averages are shown.

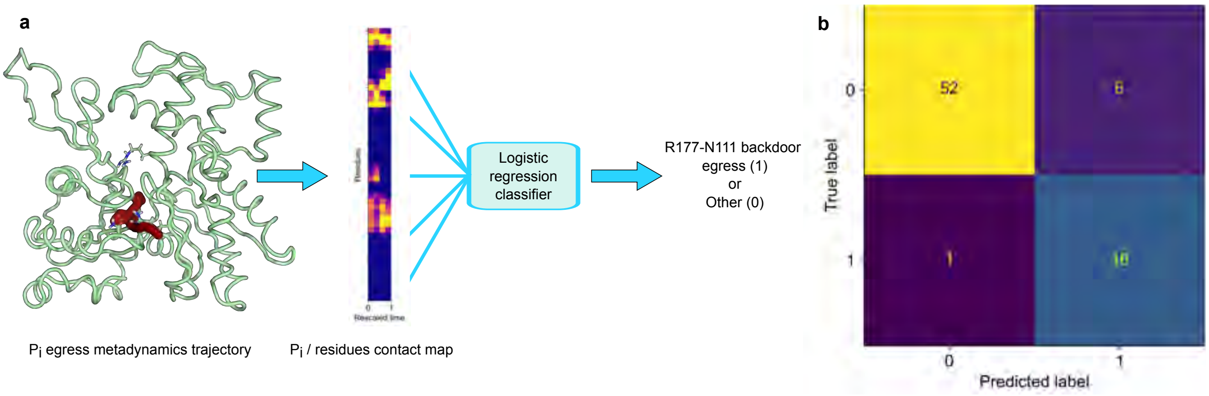


**Supplementary Fig. 13 Classifying $P_i$ egress trajectories by machine learning. a.** Machine learning strategy for detection of $P_i$ egress through the R177-N111 backdoor. **b.** Confusion matrix of the logistic regression classifier on the test set.
